# The transcriptional and translational landscape of equine torovirus

**DOI:** 10.1101/296996

**Authors:** Hazel Stewart, Katherine Brown, Adam M. Dinan, Nerea Irigoyen, Eric J. Snijder, Andrew E. Firth

**Affiliations:** Division of Virology, Department of Pathology, University of Cambridge, Cambridge, United Kingdom.; Molecular Virology Laboratory, Department of Medical Microbiology, Leiden University Medical Center, Leiden, The Netherlands.

## Abstract

The genus *Torovirus* (subfamily *Torovirinae*, family *Coronaviridae*, order *Nidovirales*) encompasses a range of species that infect domestic ungulates including cattle, sheep, goats, pigs and horses, causing an acute self-limiting gastroenteritis. Using the prototype species equine torovirus (EToV) we performed parallel RNA sequencing (RNA-seq) and ribosome profiling (Ribo-seq) to analyse the relative expression levels of the known torovirus proteins and transcripts, chimaeric sequences produced via discontinuous RNA synthesis (a characteristic of the nidovirus replication cycle) and changes in host transcription and translation as a result of EToV infection. RNA sequencing confirmed that EToV utilises a unique combination of discontinuous and non-discontinuous RNA synthesis to produce its subgenomic RNAs; indeed, we identified transcripts arising from both mechanisms that would result in sgRNAs encoding the nucleocapsid. Our ribosome profiling analysis revealed that ribosomes efficiently translate two novel CUG-initiated ORFs, located within the so-called 5’ UTR. We have termed the resulting proteins U1 and U2. Comparative genomic analysis confirmed that these ORFs are conserved across all available torovirus sequences and the inferred amino acid sequences are subject to purifying selection, indicating that U1 and U2 are functionally relevant. This study provides the first high-resolution analysis of transcription and translation in this neglected group of livestock pathogens.

**Importance:** Toroviruses infect cattle, goats, pigs and horses worldwide and can cause gastrointestinal disease. There is no treatment or vaccine and their ability to spill over into humans has not been assessed. These viruses are related to important human pathogens including severe acute respiratory syndrome (SARS) coronavirus and they share some common features, however the mechanism that they use to produce subgenomic RNA molecules differs. Here we performed deep sequencing to determine how equine torovirus produces subgenomic RNAs. In doing so, we also identified two previously unknown open reading frames “hidden” within the genome. Together these results highlight the similarities and differences between this domestic animal virus and related pathogens of humans and livestock.

## Introduction

The order *Nidovirales* currently contains four families of positive-sense, single-stranded RNA viruses: the *Coronaviridae*, *Arteriviridae*, *Roniviridae* and *Mesoniviridae* (1). Their grouping into the one taxonomic order is based upon replicase protein conservation, genome organisation and replication strategy. However these viral families are nonetheless very diverse with respect to their virion structure, host range, pathogenic potential and genome size.

The genus *Torovirus* (family *Coronaviridae*, subfamily *Torovirinae*) encompasses a range of species with worldwide distribution that infect domestic ungulates including cattle, goats, sheep, pigs and horses, causing an acute self-limiting gastroenteritis. Approximately 55 % of cattle within the United Kingdom are seropositive for bovine torovirus and this pathogen represents a significant burden to the industry (2, 3). Similarly porcine torovirus is endemic in Europe and causes disease in production herds (4–6). Despite this, limited research has been conducted upon these pathogens and neither specific antiviral treatments nor vaccines are available. The prevalence of toroviruses in non-domestic reservoirs and potential for cross-species transmission has not been assessed, although they are known to undergo recombination events (7). The extensive research conducted upon the related coronaviruses would not necessarily be relevant in the event of an emerging torovirus infection, due to the divergent nature of these viruses.

The genomes of *Nidovirales* are positive-sense, polycistronic RNAs. One of the hallmarks of this virus order is the utilisation of an unusual transcription mechanism to express the genes encoding structural and accessory proteins, which reside downstream of the large replicase open reading frames (ORFs) 1a and 1b (Figure 1). These proteins are typically translated from a nested set of 3’ coterminal subgenomic mRNAs (sg mRNAs). Although, with the exception of the smallest species, these sgRNAs are structurally polycistronic, translation is normally limited to the 5’ ORF of each mRNA. Studies of coronaviruses and arteriviruses have revealed that they produce negative-sense subgenome-sized RNAs via a mechanism of “discontinuous” extension (8). This process resembles homology-assisted copy-choice recombination (9) and requires the presence of multiple copies of a species-specific short motif, the transcription regulatory sequence (TRS). TRS motifs are located immediately upstream of the structural protein ORFs (body TRSs) and within the 5’ UTR (leader TRS).

**Figure 1.**
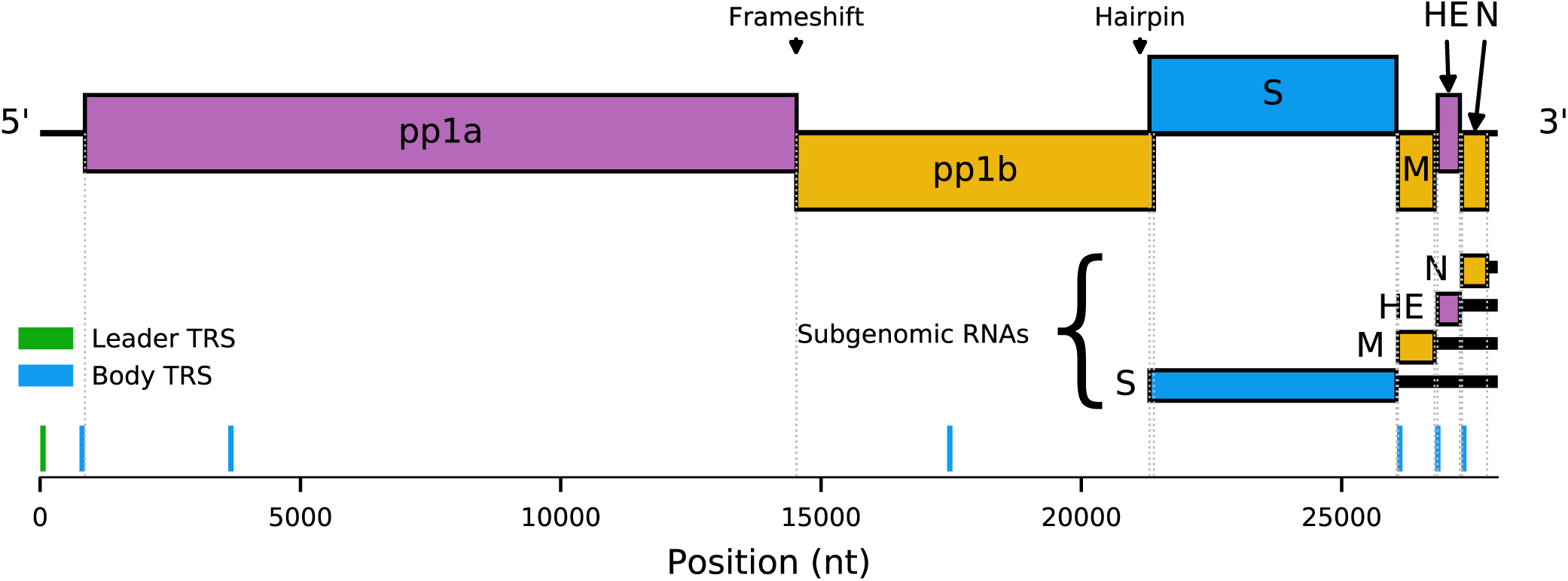
Schematic of the equine torovirus genome (EToV). Open reading frames (ORFs) are coloured according to their respective reading frames (pink: phase 0 yellow: phase −1; blue: phase +1). Polyproteins pp1a and pp1ab are translated from genomic RNA, with pp1ab generated via −1 programmed ribosomal frameshifting. Structural proteins are translated from a series of subgenomic RNAs. Untranslated regions of subgenomic RNAs are represented by black bars. The leader transcription regulatory sequence (TRS) (green) and putative body TRSs (blue) are displayed below the viral genome. The frameshift site and a putative RNA hairpin involved in S sgRNA synthesis are indicated above the genome.

Negative strand RNA synthesis initiates at the 3’ end of the positive-sense viral genome. When the RNA-dependent RNA polymerase (RdRp) has copied a TRS sequence, a translocation event may occur during which the anti-TRS at the 3’ end of the nascent strand basepairs with the leader TRS within the 5’ UTR. Transcription reinitiates and continues to the 5’ end of the genomic template. The resulting “anti-leader” sequence that is added ranges from 55 – 92 nt in coronaviruses to ~200 nt in arteriviruses. These negative-sense transcripts are therefore 5’- and 3’-coterminal with the full length negative RNA strand and are identifiable as chimaeras with distinct leader-body junctions. The anti-leader sequence in each of the negative-sense templates then functions as a promoter, to drive synthesis of a mirror set of positive-sense sgRNAs that are translated to produce the structural proteins.

However not all details of the mechanism outlined above are wholly conserved across the *Nidovirales*. Specifically, the two sg mRNAs of roniviruses (pathogens of shrimp) do not possess conserved 5’ leader sequences, indicative of the lack of a discontinuous step during their production (10). Despite the presence of a conserved body TRS in each sg mRNA, an equivalent leader TRS is not readily identifiable in the 5’ UTR. It may therefore be reasoned that the ronivirus body TRSs stimulate termination of RNA synthesis without RdRp translocation and reinitiation. Mesoniviruses (a branch of *Nidovirales* recently identified in insects) are thought to produce two major sgRNAs possessing leader sequences of different lengths, indicating the nidoviral mechanism for discontinuous RNA synthesis may allow two very different leader/body TRS pairs to be utilised in a single viral species (11).

Toroviruses appear to represent a nidovirus subgroup with a remarkably flexible transcription strategy: equine torovirus (EToV) possesses a leader TRS-like sequence (CUUUAGA) but it is only involved in the synthesis of the mRNA used for expression of the spike (S) protein gene (12). Despite similarities to the corona- and arteriviral mechanism, the preceding leader sequence incorporated into this mRNA is merely 6 nt in length (ACGUAU). Additionally, this case is unusual in that the translocation event is thought to be prompted by an RNA structure - a predicted RNA hairpin upstream of the S protein gene, rather than a body TRS (12). Body TRSs are located upstream of the three remaining structural protein genes, yet a non-discontinuous mechanism is utilised for their production, as is the case for roniviruses. As a result, the sg mRNAs for membrane (M), nucleocapsid (N) and haemagglutinin-esterase (HE) do not normally possess a conserved 5’ leader sequence; they each possess a variable and unique extended version of the TRS at their 5’ end. It is clear there is significant difference between how the various *Nidovirales* families synthesise their sgRNAs.

Here we describe the first high-resolution analysis of viral transcription during infection by EToV, which is one of the few toroviruses that can be propagated in cell culture (13, 14). RNA sequencing (RNA-seq) confirmed previous reports that EToV utilises a unique combination of both discontinuous and non-discontinuous RNA synthesis to generate its repertoire of sgRNAs. Strikingly, we also identified a small proportion of chimaeric transcripts spanning from the leader to the body TRS of the N protein gene, indicating that discontinuous and non-discontinuous mechanisms compete in this location. We also identified numerous locations across the genome where non-canonical RdRp translocation occurs, leading to a vast array of (presumably mostly non-functional) chimaeric transcripts.

Ribosome profiling (Ribo-seq) conducted in tandem with the RNA-seq indicated ribosomes were actively translating within the so-called 5’ UTR. Further analysis confirmed the existence of two novel ORFs in this region, which are conserved in all torovirus genome sequences analysed to date. The specific function(s) of these proteins will be the topic of future work. Together, these results provide an overview of the transcriptional and translational events that accompany infection by this wide-ranging pathogen.

## Results

### Tandem RNA-seq and Ribo-seq of EToV infected cells

We conducted tandem RNA-seq and Ribo-seq of EToV infected equine dermal (ED) cells. Two biological replicates of virus-infected and mock-infected cells were analysed, generating 25 to 53 million reads per sample. For RNA-seq, 77-92 % of reads mapped to the host genome, of which a mean of 1.5 % mapped to rRNA, 19 % to mRNA, 32 % to ncRNA and 47 % elsewhere in the genome. For Ribo-seq, 46-60 % of reads mapped to the host genome, of which a mean of 56 % mapped to rRNA, 13 % to mRNA, 4.9 % to ncRNA and 26 % elsewhere in the genome (Supplementary Table 1). 1.3 % and 2.3 % of reads mapped to the virus genome in the two EToV-infected RNA-seq replicates and 0.41 % and 0.21 % in the two virus-infected Ribo-seq replicates.

The viral genome was assembled *de novo* from RNA-seq reads and confirmed as EToV, Berne isolate. A single 27694-nt contig was assembled representing almost the entire viral genome. Only 18 nt at the 5’ terminus and 300 nt at the 3’ terminus of this contig failed to assemble automatically; however these regions were clearly covered by reads consistent with the reference sequence on inspection and so were added manually to the consensus sequence. Four single nucleotide changes were present in all reads but not the reference sequence compiled from previous sequencing data, at positions 18078 (ORF 1b, C > U), 21429 (ORF S, A > U), 21814 (ORF S, C > A) and 25596 (ORF S, C > U). The full-length virus sequence has been deposited in GenBank (Accession MG996765).

The distribution of reads on the virus genome and the phasing of these reads are shown in Figure 2. There was good coverage across the viral genome for both RNA-seq and Ribo-seq. The Ribo-seq/RNA-seq ratio along the genome was calculated (Figure 2C) to estimate translation efficiency (note that this simple estimate is naive since it does not account for the fact that the genomic RNA and different sgRNA species overlap one another). Ribo-seq density, RNA-seq density and translational efficiency were also calculated separately for each ORF (Figure 3), based on the density of Ribo-seq reads in each ORF divided by the density of the RNA-seq reads for either the same region (for subgenomic RNAs) or the region of the genome which does not overlap the subgenomic RNAs (for genomic RNA). RNA-seq density was adjusted based on the “decumulation” methodology described previously (15) (see Materials and Methods) to account for the fact that not all of the RNA-seq density in the 3’ ORFs derives from transcripts from which the ORFs can be expressed. Ribo-seq coverage is much higher towards the 3’ end of the genome, particularly across the M and N genes, reflecting the translation of abundant subgenomic RNAs in this region (Figure 2, Figure 3). ORFs 1a and 1b contain a considerably lower density of Ribo-seq reads. The relatively low translation efficiencies calculated for ORFs 1a and 1b may be partly due to some gRNA being packaged (or destined for packaging) and unavailable for translation but still contributing to the estimate of gRNA RNA-seq density. ORF1a has a higher Ribo-seq density and a higher translational efficiency than ORF1b, reflecting the proportion of ribosomes terminating at the ORF1a stop codon and not undergoing the −1 frameshift into ORF1b (Figure 2, Figure 3). As expected, RNA-seq density is similar across ORF1a and ORF1b, as both are present only on the full-length genomic RNA (Figure 2). The region covering the HE ORF also has low ribosomal coverage (Figure 2), which may be due to the fact that the EToV HE gene is nonfunctional due to a large deletion including the canonical AUG (16). HE is not shown in Figure 3 as the HE transcript is much less abundant than the “upstream” M transcript which makes the decumulation procedure susceptible to noise (see Irigoyen et al., 2016). Translational efficiency appears highest for the M and S subgenomic RNAs. The high RNA-seq density in the 5’ UTR may be indicative of one or more defective interfering (DI) RNAs in the sample (see below). Ribosome protected fragments (RPFs) were also identified mapping to the second half of the 5’ UTR, mostly in the +2/-1 frame with respect to ORF1a (Figure 2A).

**Figure 2.**
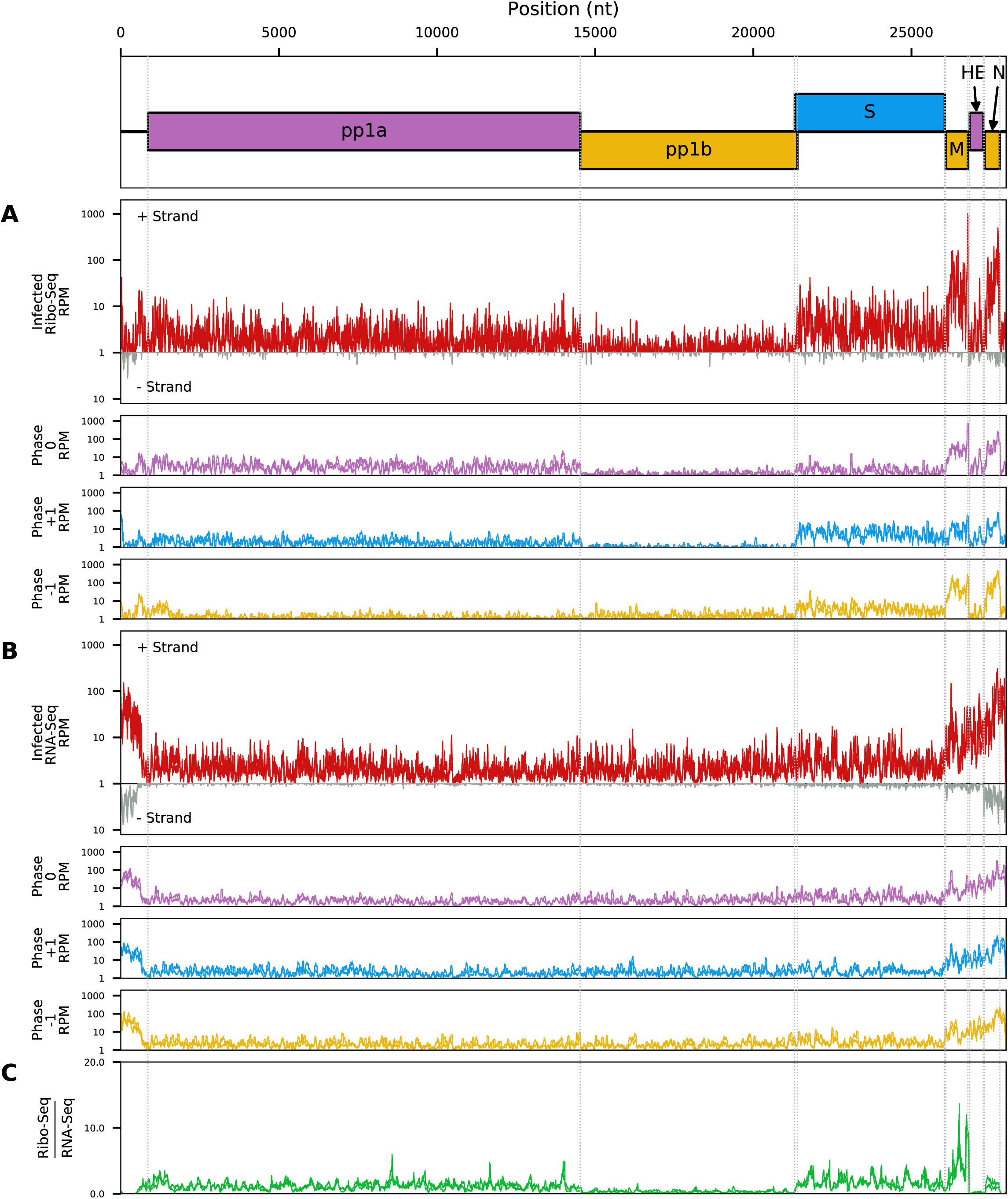
Read density of (A) Ribo-seq and (B) RNA-seq reads across the viral genome from EToV infected cells. Red lines represent total reads per million mapped reads at each position; pink: reads in phase 0; yellow: phase −1; blue: phase +1. Densities are smoothed with a 15-nt running mean filter and plotted on a log_10_(1+x) scale. Negative-sense reads (grey) are displayed below the x-axis for total reads only. Each line represents a single replicate. For Ribo-seq reads, a +12 nt offset has been applied to read 5’ end positions to map approximate P-site positions. (C) The positive sense Ribo-seq/RNA-seq ratio after applying a 100-nt running mean filter to each distribution. Each line represents one of the two paired Ribo-seq and RNA-seq replicates.

**Figure 3.**
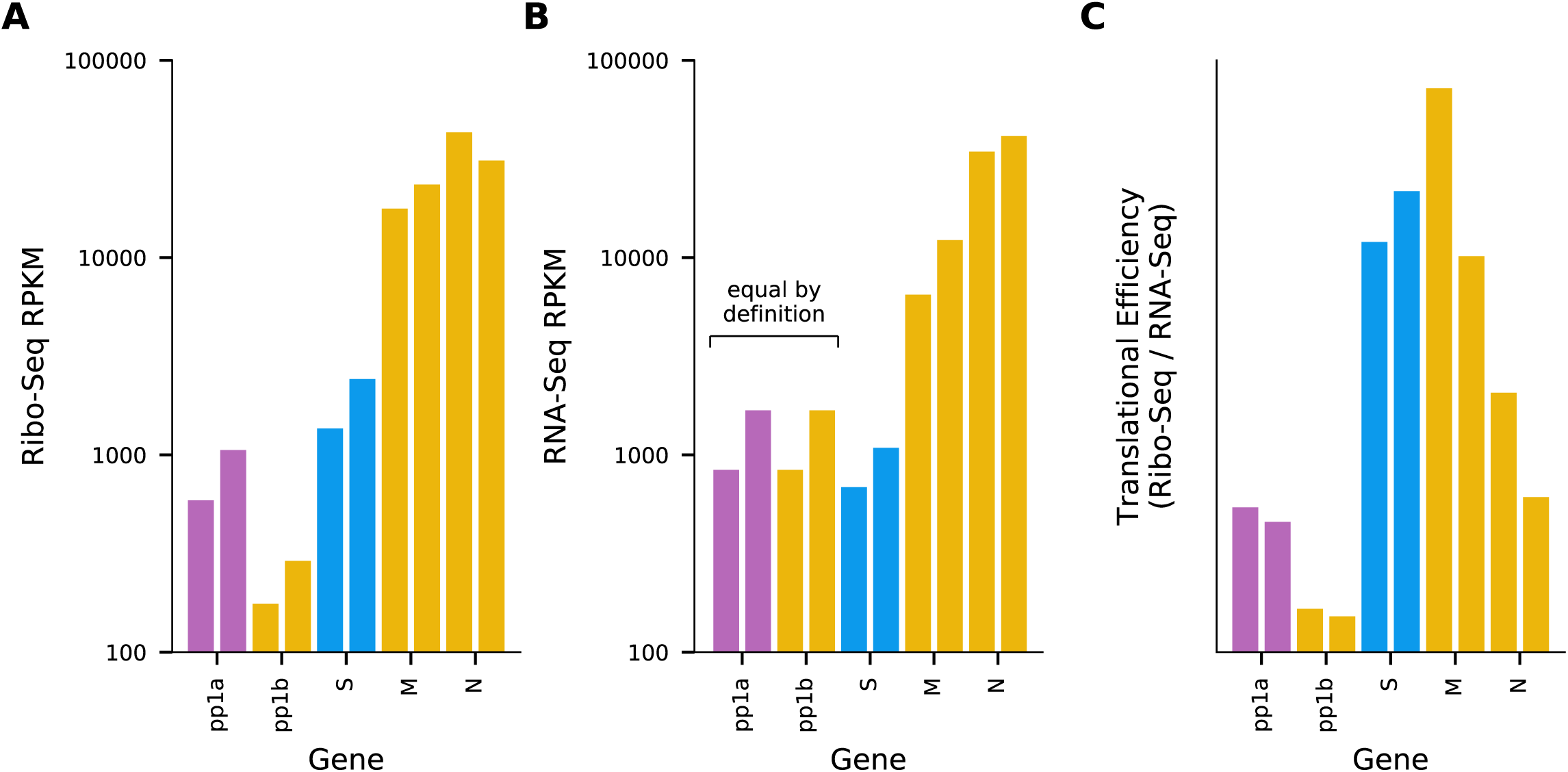
Relative gene expression levels. (A) Ribo-seq density in reads per kilobase per million mapped reads (RPKM) for each ORF in the EToV genome. For each ORF, only reads mapping in the predominant phase (i.e. mapping to first positions of codons) were included. (B) “Decumulated” RNA-seq density in RPKM for each ORF. For subgenomic RNAs, density was calculated across the regions used for Ribo-seq in A; for genomic RNAs the regions for ORF1a and ORF1b were combined, as these ORFs are both translated from gRNA. A decumulation strategy was used to correct for the fact that the measured RNA density in 3’ ORFs derives from multiple 3’-coterminal transcripts (see Materials and Methods). (C) Translation efficiency for each gene in the EToV genome, calculated as Ribo-seq density / decumulated RNA-seq density. For each ORF, the two bars represent two repeats.

To calculate the length distributions of host- and virus-mapped RPFs, we used reads mapping within coding regions. After adaptor trimming, the majority (75 %) of Ribo-seq reads were 27 – 29 nt in length, which is consistent with the expected size of mammalian ribosome footprints. As expected, the distribution of read lengths for RNA-seq was much broader, peaking between 60 and 70 nt (Supplementary Figure 1). For quality control, histograms of the 5’ end positions of host mRNA Ribo-seq and RNA-seq reads relative to initiation and termination codons were constructed (Supplementary Figures 2, 3). This confirmed we had high quality RPFs arising from host transcripts, with strong triplet periodicity (“phasing”) and very few reads mapping to 3’ UTRs. As in other datasets, a ramp effect of decreased RPF density was seen over a region of ~30 codons following initiation sites; but, unusually, in this dataset we did not observe a density peak at the initiation site itself (cf. Irigoyen et al 2016). This may be due to the flash freezing without cycloheximide pretreatment used for these samples, as for a later cycloheximide-treated sample this peak is present (Supplementary Figure 2). Within coding sequences, the 5’ ends of the majority of reads from the host (65-81 %) and virus (60-75 %) mapped to the first positions of codons (Supplementary Figure 4).

The relative RPF density allowed us to estimate the efficiency of ribosomal frameshifting in the context of virus infection. After translating ORF1a, a proportion of ribosomes undergo a −1 ribosomal frameshift to translate ORF1b (17). This is (presumably) required to produce a specific ratio of pp1a to pp1ab, thereby controlling the ratio of RNA-synthesing enzymes such as RdRp and helicase to other components of the replicase complex, including the proteinases and trans-membrane subunits encoded in ORF1a. The ORF1a/1b −1 ribosomal frameshifting event is stimulated by a pseudoknot structure 3’-adjacent to the U_UUA_AAC slippery heptanucleotide frameshift site. The efficiency of −1 ribosomal frameshifting (measured by dividing the mean RPF density in ORF1b by the mean density in ORF1a) was estimated to be 29.9 % for replicate one and 27.5 % for replicate 2, which is in accordance with the rates measured previously outside of the context of virus infection (20 – 30 %) (17).

### RNA sequencing indicates both discontinuous and non-discontinuous mechanisms are utilised for N protein gene sgRNA synthesis

RNA sequencing reads that did not map to either the viral genome or host databases were analysed for containing potential viral chimaeric junctions, indicative of leader-to-body joining during discontinuous sgRNA synthesis (Figure 4). Relative abundances were calculated by normalising read counts to the number of non-chimaeric reads spanning each junction. Between the two replicates combined, 8330 reads were identified as chimaeras, mapping to 2837 putative junction sites. Of these, 213 were considered to be highly supported by the data, either due to being identified in at least 10 chimaeric reads or containing the full 5’ leader and TRS sequence. Adjacent donor or acceptor sites were then merged (see Materials and Methods), leaving 70 unique junctions (Figure 4).

**Figure 4.**
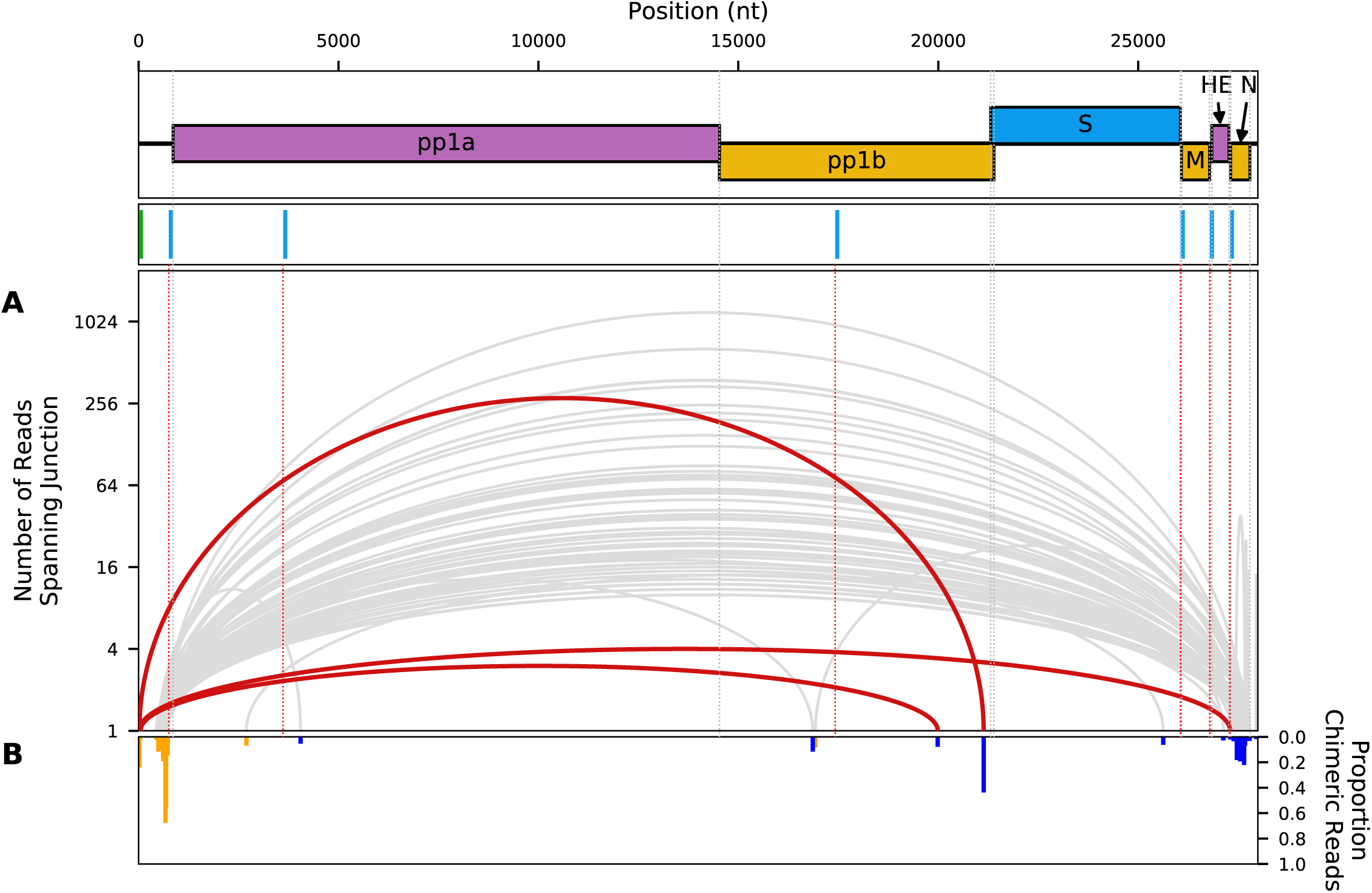
Analysis of chimaeric viral reads. (A) Sashimi plot showing junctions in the EToV genome across which chimaeric RNA-seq reads were identified in EToV infected, non-drug treated samples. Chimaeric reads were defined as reads for which the intact read could not be mapped but for which the 5’ and 3’ ends could be uniquely mapped to non-contiguous regions of the EToV genome. Junctions that were either covered by at least 10 chimaeric reads (grey) and/or for which the 5’ section of the read contained the full 5’ leader sequence and leader TRS (red) were identified and adjacent positions merged. These junctions are shown as curved lines connecting the position of the 3’ end of the 5’ mapped segment of the read and the 5’ end of the 3’ mapped segment of the read. The apical height of each curved line shows the absolute number of reads spanning this junction on a log_10_(1+x) scale. (B) Inverted bar chart showing, for the 5’ (orange) and 3’ (blue) breakpoints for each junction, the number of chimaeric reads as a fraction of the total number of chimaeric and non-chimaeric reads at each site.

Three chimaeric junctions were identified where the first nucleotide of the corresponding read mapped to the first nucleotide of the viral genome. Of these, one junction was consistent with the previously characterised sgRNA produced via discontinuous RNA synthesis encoding the S gene (280 reads, or 3 % of total chimaeric reads) (12). These reads spanned the entire leader-body junction of the S gene, possessing 14 - 18 nt of the 5’ UTR (i.e. the actual 5’-derived sequence is at least 14 nt, ACGUAUCUUUAGAA, comprising the so-called 6-nt leader, the leader TRS CUUUAGA, and an additional A), followed by the stretch of ORF1b just upstream of the S gene. A second set of transcripts containing 5’ leader sequence was identified by four unique reads starting with the 5’ leader (ACGUAU) and TRS sequence (CUUUAGA), where the remainder of the read mapped to the start of the N gene. This indicates that, contrary to previous reports, low levels of discontinuous RNA synthesis are used during production of the N gene negative-strand RNA. The final chimaera which included the 6 nt leader was represented by three reads. These reads included 44 - 46 nt of the 5’ UTR (i.e., significantly more than the predicted leader-TRS) followed by a sequence mapping to position 19987-19989 which is within ORF1b.

A substantial number of additional chimaeric reads were identified, indicative of non-TRS-driven cases of discontinuous RNA synthesis, although formally it is possible that some of these are template-switching artefacts introduced during library preparation and/or sequencing. Additionally, a large number of reads spanning from the 5’ UTR to either within the N protein gene or the 3’ UTR were identified. Indeed, the only junction represented by over 1000 reads spanned nucleotides 673 to 27649; similarly the second most commonly identified junction spanned 687 to 27550 (642 reads). If chimaeric reads were predominantly a sequencing artefact, the abundance of any particular chimaera would be approximately proportional to the product of the abundances of the sequences from which the 5’ and 3’ ends of the chimaera are derived (with some variation due to sequence-specific biases), and thus a high density of chimaeras would be expected to fall entirely within the N transcript. In contrast, most of the observed chimaeric reads were between N and the 5’ UTR. The relative paucity of reads mapping to generic locations in the ORF1ab region also argues against the majority of chimaeras being simply artefactual. The 5’ UTR preference may be due to genome circularisation during negative-sense synthesis as has been proposed for coronaviruses (18). Alternatively these may derive from autonomously replicating defective interfering RNAs, rather than multiple independent RNA translocation and reinitiation events. Such defective interfering RNAs have been extensively analysed previously and are a common complication of EToV studies (19). Consistent with the high level of 5’UTR:N chimaeric sequences, there was high RNA-seq density throughout much of the 5’ UTR, with the 3’ extent of the region of high density coinciding approximately with the region to which a large number of the chimaeric 5’ ends mapped (Figure 2, Figure 4).

### Gene expression analysis indicates multiple pathways are perturbed by EToV infection

The RNA-seq data were analysed to identify genes that were differentially expressed between virus-infected and mock-infected ED cells. We identified 61 genes that were upregulated in virus-infected cells; amongst which eight gene ontology (GO) terms were overrepresented, mostly related to the nucleosome or immune responses (Figure 5). We found 24 genes that were downregulated in infected cells, amongst which four GO terms were overrepresented, two of which were related to the ribosome. We also analysed differential translational efficiency (based on the RPF to mRNA ratio) between mock- and virus-infected cells. We identified 22 genes that were translated more efficiently in infected cells; GO analysis indicated that these genes tend to encode proteins that are involved in RNA binding. Only two genes were found to be translated less efficiently in infected cells compared to mock (Supplementary Table 2 and Figure 4). Note that these analyses measure changes in individual genes relative to the global mean and do not inform on global changes in host transcription or translation as a result of virus infection.

**Figure 5.**
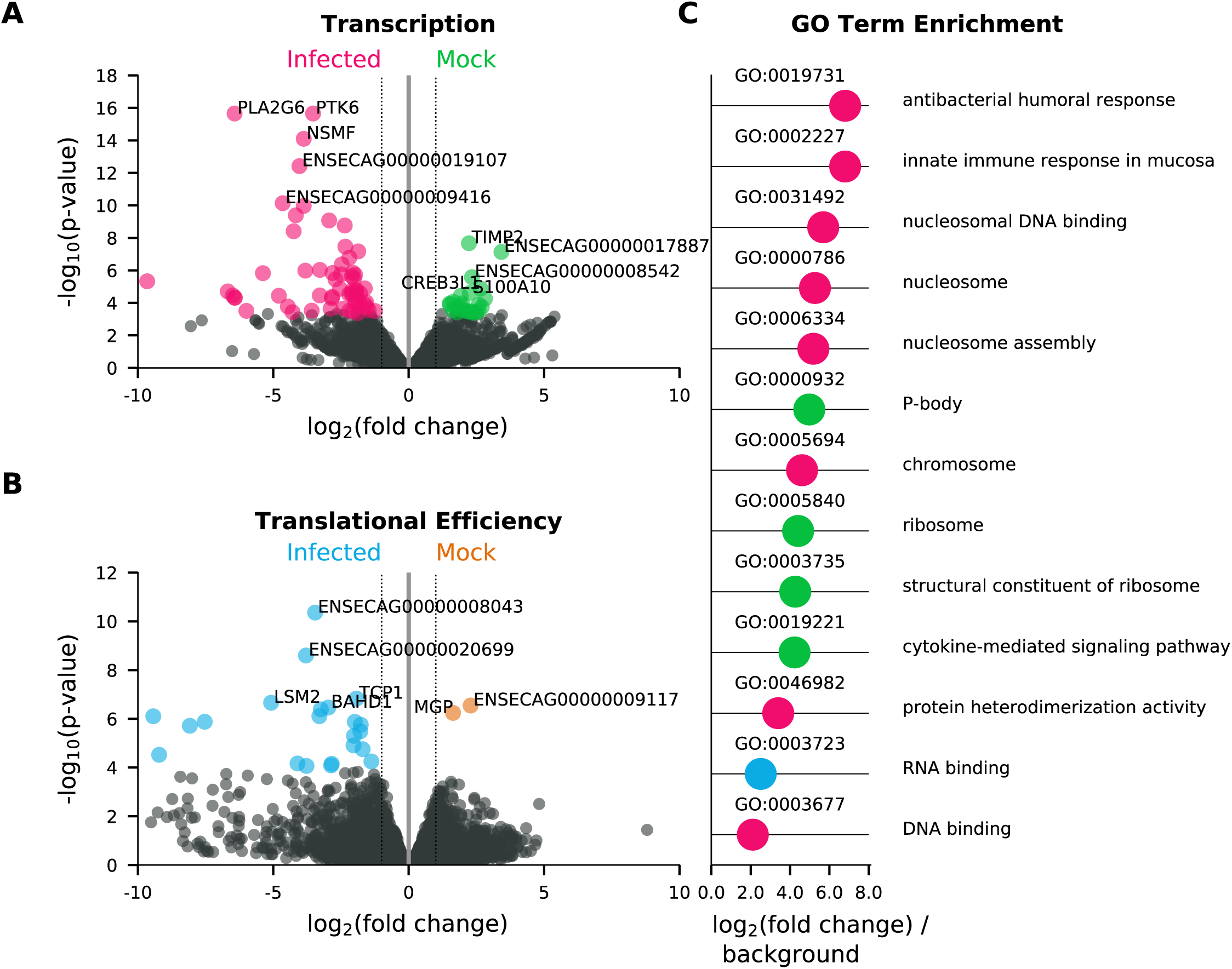
Volcano plots showing the results of (A) differential transcription analysis performed using DESeq2 (64) and (B) differential translation efficiency analysis performed using Xtail (68), between cells infected with EToV (infected) and uninfected cells (mock). Genes which were expressed at significantly higher levels (FDR ≤ 0.05 and absolute log_2_(fold change) ≥ 1) in infected cells are highlighted in pink (transcription, A) and blue (translational efficiency, B). Genes which were expressed at significantly higher levels in mock infected cells are highlighted in green (transcription, A) and orange (translational efficiency, B). The five most significant genes in each category are labelled with the gene symbol where available and otherwise with the Ensembl gene ID. (C) Absolute log_2_(fold change) for all gene ontology (GO) terms which were significantly overrepresented compared to a background of all horse protein-coding genes for genes significantly more transcribed in infected cells (pink), genes significantly more efficiently translated in infected cells (blue) and genes significantly more transcribed in mock cells (green). No terms were identified for genes significantly more efficiently translated in mock cells.

### Two additional proteins are translated from 5’ CUG-initiated ORFs

Our initial dataset indicated an excess density of ribosomes translating within the +2/-1 frame upstream of ORF1a and overlapping the 5’ end of ORF1a (Figure 2A). To further investigate this, we repeated the ribosome profiling using infected cells treated with translation inhibitors prior to flash freezing (harringtonine, HAR, and/or cycloheximide, CHX). HAR specifically arrests initiating ribosomes whilst allowing “run-off” of elongating ribosomes; conversely CHX stalls elongating ribosomes whilst allowing on-going accumulation at initiation sites. Our quality control analysis confirmed the datasets were of similar quality to our previous experiment (Supplementary Figures 1, 2 and 4) and mapping of the RPFs provided good coverage of the EToV genome (Figure 6).

**Figure 6.**
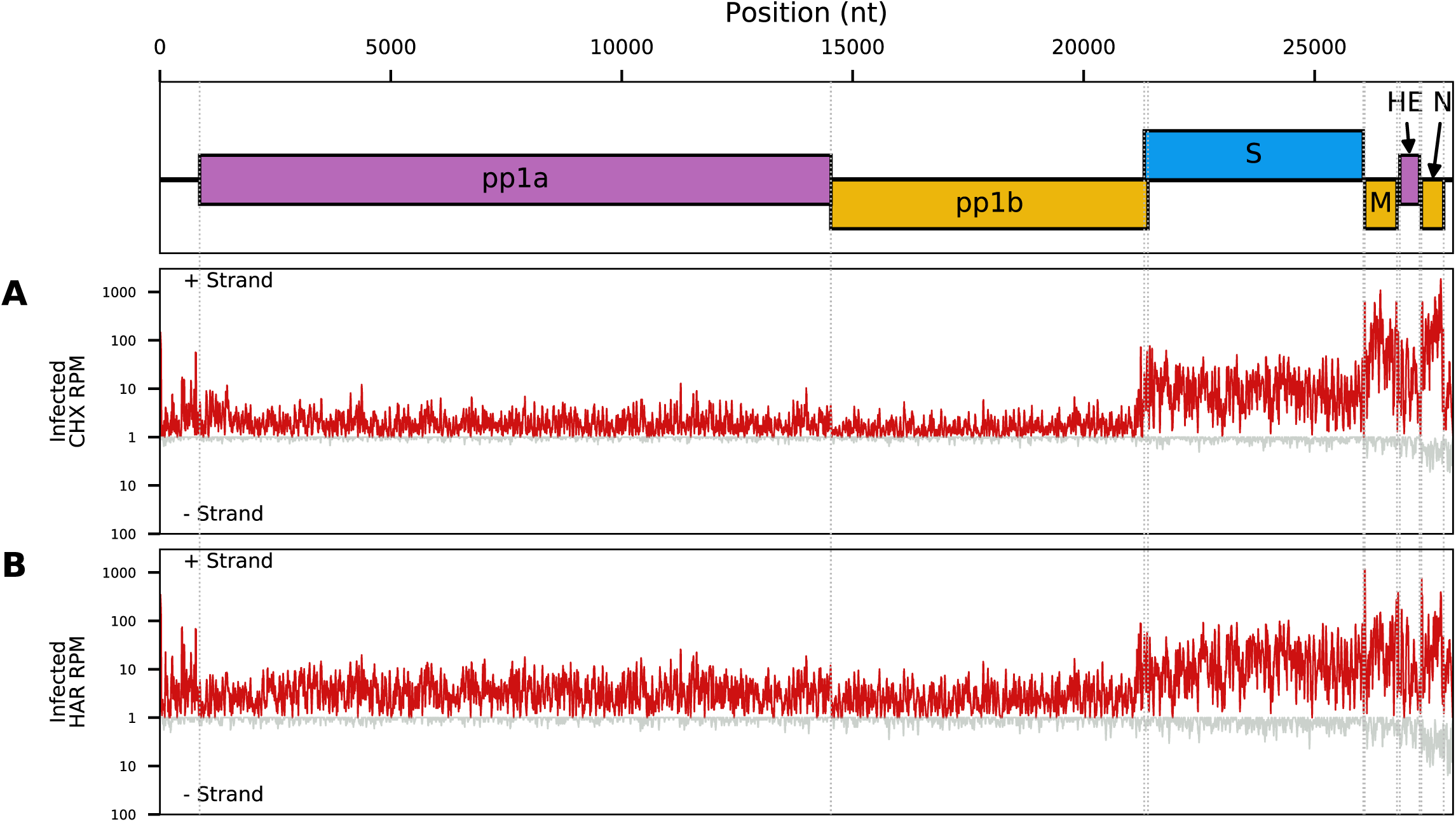
Read density of Ribo-seq reads along the viral genome for EToV infected cells pretreated with (A) cycloheximide or (B) harringtonine. Red lines represent total reads per million mapped reads (RPM) at each position. Densities are smoothed with a 15-nt running mean filter and plotted on a log_10_(1+x) scale. Negative-sense reads (grey) are displayed below the x-axis. Each line represents a single replicate. A +12 nt offset has been applied to read 5’ end positions to map approximate P-site positions.

This Ribo-seq data confirmed translation of two ORFs located within the so-called 5’ UTR and overlapping the 5’ end of ORF1a. We have termed these U1 (80 codons) and U2 (258 codon). We predict that translation of both U1 and U2 is initiated from CUG codons, as a close inspection indicated that ribosomes accumulated at these two sites (Figure 7). It must be noted that pretreatment with CHX or HAR can introduce artefacts into ribosome profiling data: CHX can lead to an excess of RPF density over ~30 codons following initiation sites when cells are stressed (15, 20). It has also been suggested that both drugs can promote upstream initiation due to scanning pre-initiation complexes stacking behind ribosomes paused at canonical initiation sites (21). However, the distance between the U1 CUG, the U2 CUG and the ORF1a initiation site, besides observation of efficient translation of U2 downstream of the ORF1a initiation site makes these artefacts unlikely to be significant confounding factors in the case of U1 and U2.

**Figure 7.**
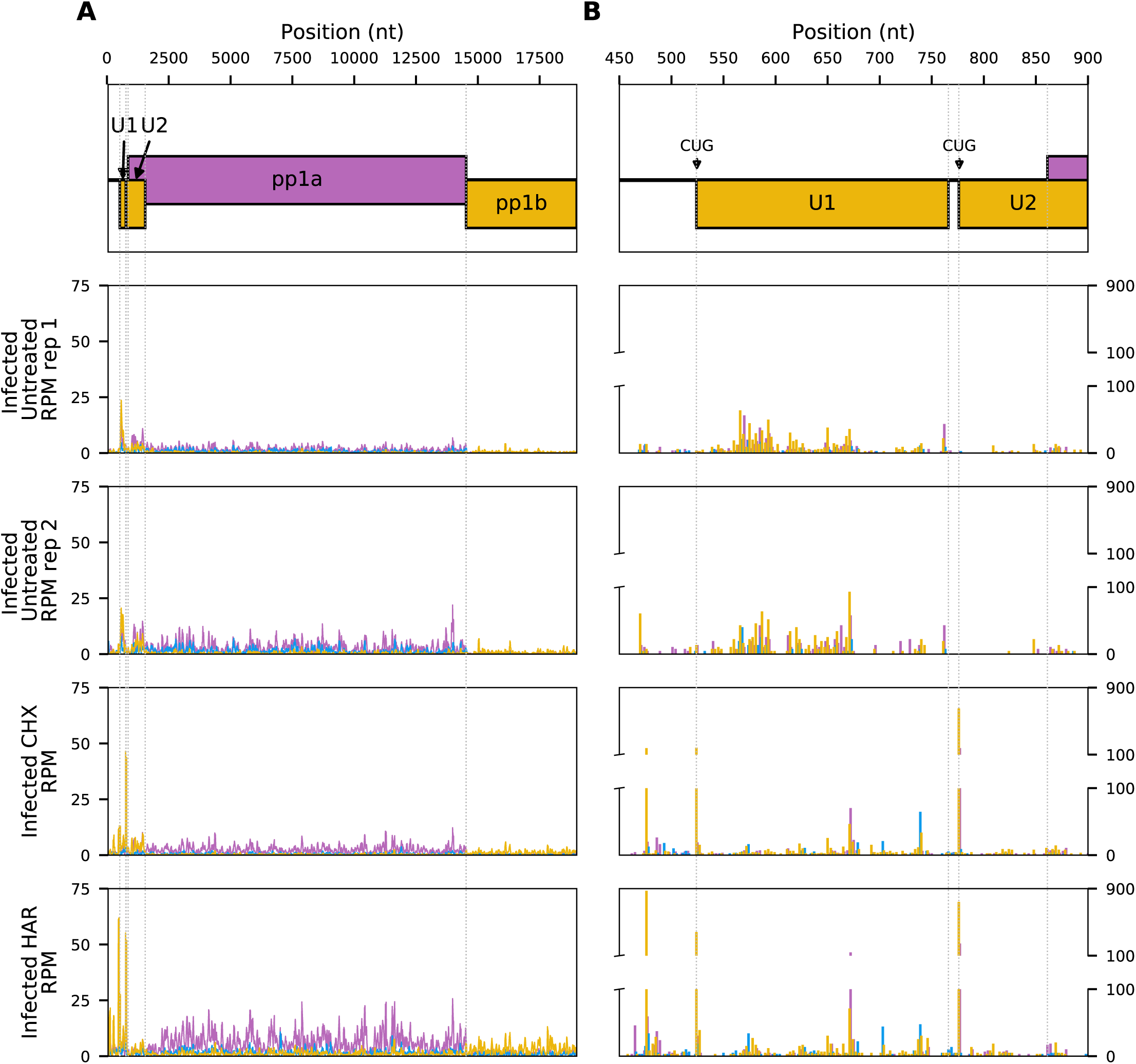
Read density of Ribo-seq reads across (A) U1, U2 and ORF1a; and (B) the U1 ORF and surrounding regions, for EToV infected cells with no drug treatment or with cycloheximide or harringtonine pretreatment. Pink: reads in phase 0; yellow: phase - 1; blue: phase +1. Graphs show total reads per million mapped reads (RPM) at each position. In (A) densities are smoothed with a 15-nt running mean filter while (B) shows the RPM counts at single-nt resolution. Each plot represents a single replicate. A +12 nt offset has been applied to read 5’ end positions to map approximate P-site positions.

Revisiting our first non-drug-treated dataset, we calculated the RPF densities and translational efficiencies within the U1 and U2 ORFs (Figure 8). U1 has a higher translational efficiency than any of the other ORFs translated from genomic RNA, whereas U2 has a translational efficiency similar to that of ORF1a.

**Figure 8.**
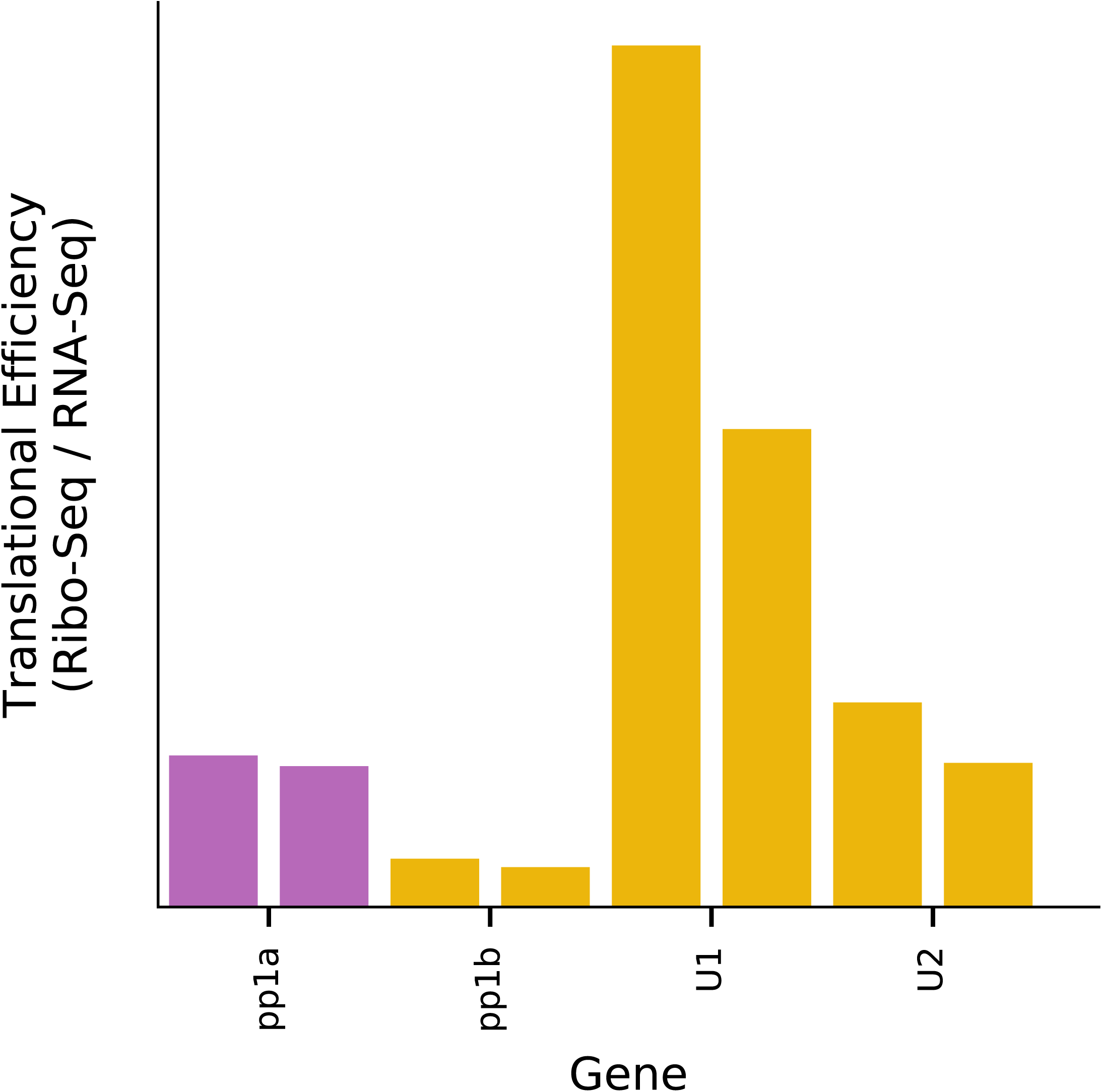
Relative translation efficiencies for U1, U2, ORF1a and ORF1b. To reduce misassignment of reads in the U2/ORF1a overlap region, for all ORFs only reads mapping in the predominant phase (i.e. mapping to first positions of codons) were included. Ribo-seq densities were divided by the ORF1ab RNA-seq densities for the corresponding paired sample. For each ORF, the two bars represent two repeats.

To assess the coding potential of U1, we calculated the ratio of non-synonymous to synonymous substitutions (dN/dS), where dN/dS < 1 indicates selection against non-synonymous substitutions which is a strong indicator that a sequence encodes a functional protein. Application of codeml (22) to a codon alignment of eight torovirus U1 nucleotide sequences resulted in a dN/dS estimate of 0.31 ± 0.08, indicating that the U1 ORF is likely to encode a functional protein. MLOGD (23) uses a principle similar to the dN/dS statistic but also accounts for conservative amino acid substitutions (i.e. similar physico-chemical properties) being more probable than non-conservative substitutions in biologically functional polypeptides. MLOGD 3-frame “sliding window” analysis of a full-genome alignment revealed a strong coding signature in the known protein-coding ORFs (as expected) and also in the U1 ORF (Figure 9).

**Figure 9.**
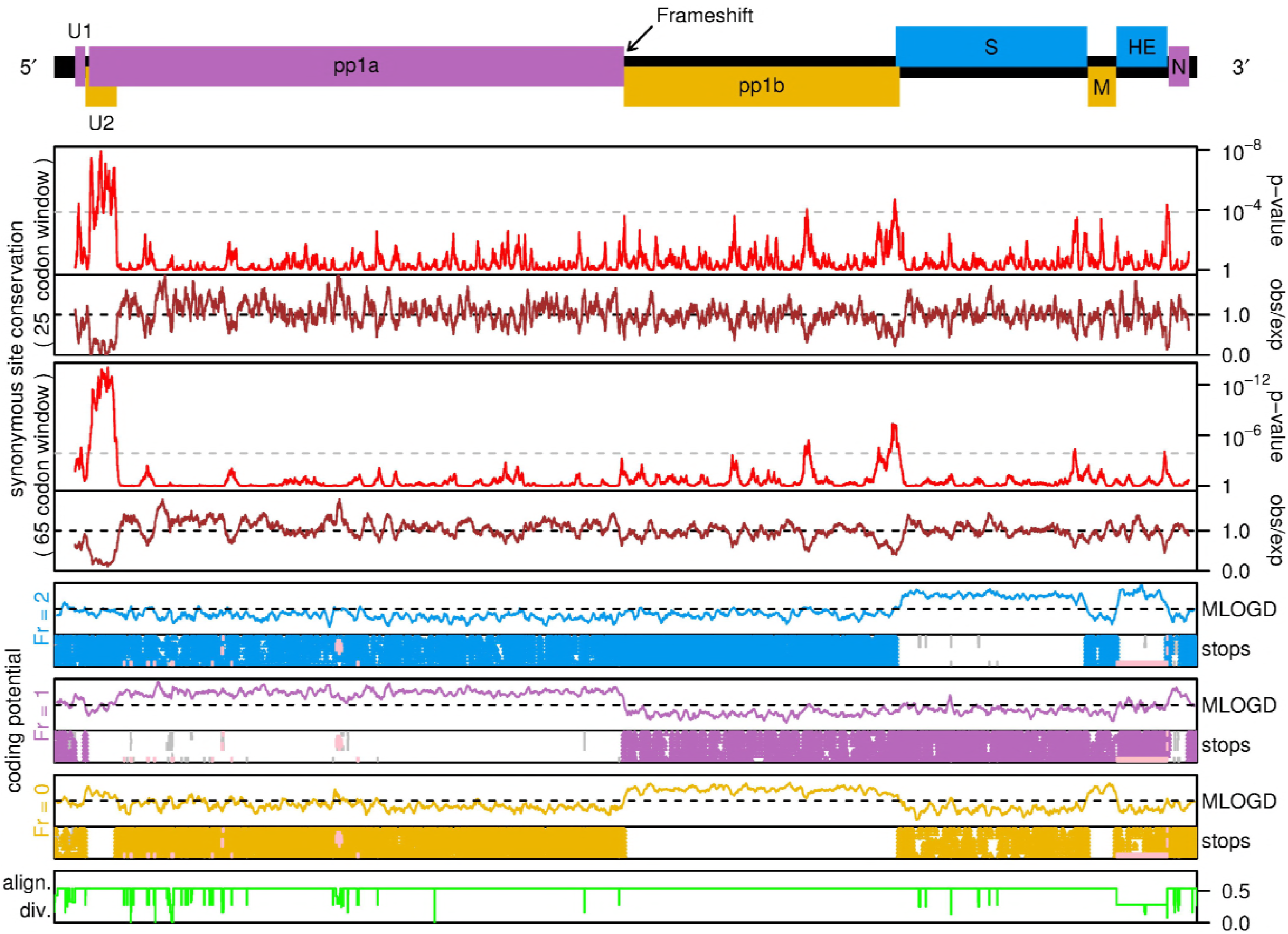
Coding potential statistics for the torovirus genome. A map of the torovirus genome is shown at top. Breda virus (AY427798.1) was used as the reference genome for this analysis since EToV has a deletion in the HE gene. In Breda virus, U1 is in-frame with ORF1a due to a 2-nt insertion relative to EToV in the short non-coding region between U1 and U2. The next four panels show an analysis of synonymous site conservation in the concatenated coding ORFs (with the reading frame of the longer ORF being used wherever two ORFs overlap). Red lines show the probability that the degree of conservation within a given window (25- or 65-codons as indicated) could be obtained under a null model of neutral evolution at synonymous sites, whereas brown lines depict the absolute amount of conservation as represented by the ratio of the observed number of substitutions within a given window to the number expected under the null model. Greatly enhanced synonymous site conservation is seen in the region of ORF1a that is overlapped by the U2 ORF. The next three panels show MLOGD coding potential scores and stop codon plots for each of the three reading frames. The positions of stop codons are shown for each of the eight torovirus sequences mapped onto the Breda virus reference sequence coordinates. Note the conserved absence of stop codons in the U1 and U2 ORFs. MLOGD was applied in a 40-codon sliding-window (5-codon step size). Positive scores indicate that the sequence is likely to be coding in the given reading frame. Note the positive scores within the U1 and U2 ORFs besides the previously known ORFs. The bottom panel (green line) indicates the total amount of phylogenetic divergence contributing to the analyses at each alignment position (regions containing alignment gaps have reduced summed divergence leading to reduced statistical power). Pink regions in the stop codon plots (e.g. EToV sequence in the HE region) indicate regions excluded from the analyses due to poor or locally out-of-frame mapping to the Breda reference sequence (see Firth, 2014 for details).

We previously predicted the existence of U2 via an analysis of coding potential and synonymous site conservation across the two torovirus genomes available at that time (24). Six additional torovirus genome sequences have now become available. We therefore extended the bioinformatics analysis using all eight currently available torovirus genome sequences (Figure 9). Since the U2 ORF overlaps ORF1a, leading to constraint on dS, the dN/dS analysis is not appropriate for U2. MLOGD analysis indicated that the U2 ORF has a higher coding potential than the corresponding part of ORF1a (Figure 9). Overlapping genes are thought mainly to evolve through “overprinting” of an ancestral gene by the *de novo* gene (25). The *de novo* gene product is often an accessory protein and often disordered (26). Interestingly, the fragment of pp1a encoded by the region of ORF1a that is overlapped by U2 has no tblastn (27) nor HHpred (28) homologues outside of the *Torovirus* genus. Thus, it is unclear which of U2 and the N-terminal domain of pp1a is ancestral. To provide further comparative genomic evidence for the functionality of U2, we used synplot2 to assess conservation at synonymous sites in the ORF1a reading frame, since overlapping functional elements are expected to place extra constraints on synonymous site evolution (29). Consistent with the earlier 2-sequence analysis (24), synplot2 revealed greatly enhanced ORF1a-frame synonymous-site conservation in a region coinciding precisely with the conserved absence of stop codons that defines the U2 ORF (Figure 9), with the mean rate of synonymous substitutions in that region being 0.20 of the genome average. Summed over the 230-codon overlap region, the probability that the observed level of conservation would occur by chance is *p* = 6.5 × 10^−40^.

Both U1 and U2 are conserved in all eight torovirus sequences with no variation in length or initiation or termination position (Supplementary Figure 5). In all sequences, U1 and U2 begin with a CUG codon in a strong initiation context (‘A’ at −3 for U1, and ‘A’ at −3 and ‘G’ at +4 for U2) (30). The U1 protein is predicted to contain two central transmembrane domains and has a C-terminus containing many charged amino acids. The U2 protein is predicted to form alternating α helix and antiparallel β sheet domains, however no structural homologs were found through searches of public databases (31–33). Their function(s) will be the topic of future work.

## Discussion

### RNA-seq reveals the complexity of torovirus transcription mechanisms

The factors influencing which transcriptional mechanism is utilised for the synthesis of each sgRNA during torovirus replication have not been elucidated. The EToV genome contains seven occurrences of the canonical TRS motif (CUUUAGA): within the 5’ UTR (leader TRS), the end of U1, central ORF1a, central ORF1b, and immediately before the M, HE and N ORFs (Figure 1). Consistent with experimental evidence (12), we did not identify any chimaeric transcripts encompassing the body TRS of M or HE, or those within ORF1b or ORF1a. It appears that these sites do not stimulate interruption of negative strand RNA synthesis followed by subsequent re-pairing and reinitiation. The nucleotides flanking the N, M and HE TRSs are semi-conserved (Supplementary Figure 6) and it has been suggested previously that the motif definition should be extended to cACN_3-4_CUUUAGA to reflect this (34). It is likely that these flanking nucleotides contribute to the degree of utilisation.

For the S gene, the chimaeric junction occurs within the run of uridines 3’-adjacent to the hairpin (Figure S6I). Our results lend support to the hypothesis suggested previously that a short conserved RNA hairpin, 174 nt upstream of the AUG start codon of the EToV S protein gene, mediates discontinuous extension of negative strand RNA synthesis to produce this sgRNA (12) (Supplementary Figure 6). The predicted hairpin structure was not present in S gene chimaeric reads, indicating that translocation may indeed be prompted by the RdRp encountering a physical block after synthesising the reverse complement of the S ORF. This is in contrast to the coronaviral and arteriviral mechanism, wherein RNA structures are insufficient and an accompanying body TRS is required to act as a transcriptional attenuation signal, prompting translocation and re-pairing of the nascent RNA. We cannot unambiguously identify which nucleotides are templated before or after the translocation event, as a GUUU sequence maps to genomic RNA on either side of the breakpoint.

The leader-TRS chimaeric reads mapping to the N protein gene initially appear consistent with the coronaviral and arteriviral mechanism of TRS-driven discontinuous RNA synthesis. However close inspection indicated that the homologous motif mediating copy-choice recombination-like translocation and re-pairing of RNA strands was actually a short AGAA sequence, not the true TRS (tetranucleotides underlined in Figures S6A and S6G). This would result in the nascent anti-TRS mispairing with the leader TRS; two nucleotides are “skipped” once reinitiation occurs. This may explain why the discontinuous mechanism is utilised so rarely for this mRNA.

This leads to the suggestion that homology between any two sites may be sufficient to induce discontinuous RNA synthesis, i.e. that provided adequate sequence homology exists, the nascent RNA strand may re-pair with upstream sites within the genomic RNA regardless of the presence of a predefined TRS. This is consistent with the 5’ UTR-ORF1b chimaeric transcripts, which again revealed a particular sequence that could be templated from either region, in this case AACCUUA rather than the TRS.

If TRS sequence-specificity is not required to stimulate EToV discontinuous RNA synthesis, it is presumably constrained by alternative roles. The highly conserved nature of the canonical leader, M, HE and N TRS (CUUUAG[A/U]) across all torovirus genomes (Supplementary Figure 6) suggests it is not tolerant to mutations, however this has not been formally confirmed. Lack of conservation of the EToV U1, ORF1a and ORF1b TRS sequences is consistent with them not being functionally relevant. Our results indicate this essential nature is likely due to a role in transcriptional termination, as we did not identify a significant role of this motif in the generation of chimaeric transcripts. Conversely, the upstream region of the “extended” TRS (cACN_3-4_CUUUAGA) is tolerant to modifications, reflecting the variable nature within sequences; even when this spacer is extended to six nucleotides, transcripts are still detectable at 20 % of WT levels (34). Again, this is consistent with a role in termination rather than a requirement for re-pairing with upstream sequences. The canonical TRS sequences also presumably contribute to subgenomic promoter recognition, as the initial CAC is essential though the adenylate is the first nucleotide on all positive-strand subgenomic transcripts (34). Initiation of sgRNA transcription at AC dinucleotides is also found in the roniviruses (10). It may be that in these *Nidovirales* families, the conserved TRS is utilised primarily for signalling transcriptional termination followed by promoter recognition, and any use for discontinuous RNA synthesis is merely a byproduct of RdRp promiscuity.

The unique combination of discontinuous and non-discontinuous mechanisms within the one virus so far appears unique to the mammalian toroviruses. The one bafinivirus isolated to date (white bream virus, family *Coronaviridae*, subfamily *Torovirinae*, genus *Bafinivirus*) has an extended TRS sequence (CA[G/A]CACUAC) which is not conserved with the mammalian toroviruses analysed in this study. Bafinivirus replication produces three sgRNAs which share an identical 42-nt leader also found at the far 5’ terminus of the genome, indicating this species utilises discontinuous RNA synthesis in a manner similar to the corona- and arteriviruses (35). However there was preliminary evidence that two of the three sgRNAs exhibit diversity in their junction sites, suggesting the anti-TRS may bind to multiple sites within the 5’ leader during strand transfer, consistent with our suggestion that whilst a threshold level of homology is required this is not limited to particular primary sequences. This is reflected in the fact that the bafinivirus leader-TRS is not fully identical to the body TRSs.

It is not known which mechanism was utilised by the last common ancestor of nidovirids, and thus which represents divergence from the original model. It has been suggested that convergent evolution has resulted in the mechanism for discontinuous negative strand synthesis arising multiple times within the *Nidovirales*. Similarly, whether the initial role of the TRS motif was to merely stimulate the attenuation of RNA synthesis or to direct the discontinuous mechanism is not known. Our data suggests that transcription mechanisms in the *Nidovirales* fall into multiple categories, each requiring a distinct role of the TRS: (i) homology-driven reinitiation (canonical discontinuous RNA synthesis, as seen in coronaviruses and arteriviruses and to a low extent, EToV N protein-coding mRNAs); (ii) structure-driven discontinuous transcription (EToV S <u>protein</u> gene); and (iii) transcription termination (EToV M, HE and the majority of N protein-coding transcripts). These mechanisms all require a RdRp which is prone to translocating when even relatively short homologous sequences are present, potentially leading to a large number of irrelevant transcripts being produced (as previously observed in an arterivirus (36)) and also facilitating the production of defective interfering RNAs (34) and recombinant strains (7).

### Effects upon the host: transcriptional and translational differential expression

The differential transcription analysis indicated that infection with EToV induces increased transcription of multiple genes, the products of which are significantly more likely than random to be involved in (i) nucleosome function and DNA binding, and (ii) immune responses to infection than genes which were not differentially transcribed. Some of the identified GO categories, including cytokine signalling, innate immune responses and ribosome biogenesis have been identified in previous RNA-seq analyses of various coronaviruses (37, 38). Similarly, although differential translational analyses or proteomic studies have not been conducted upon toroviruses, some of the identified proteins have been recognised as being incorporated into nidovirid virions (for example, TCP-1 and multiple heat shock proteins within arterivirus particles) (39). Others have been identified as being upregulated upon infection with coronaviruses, such as the solute carrier family 25 members (40). Notably, both poly(C) and poly(A) binding proteins were preferentially translated in infected cells; these have been previously identified as interaction partners of arteriviral non-structural protein 1β and contribute to viral RNA replication (41). It therefore appears that torovirus infection induces a similar host response to many nidovirids.

To the best of our knowledge, this is the first analysis of differential gene expression following infection with a torovirus. It would be of interest to repeat this analysis at later time points, as a previous study found that EToV-mediated global inhibition of host protein synthesis was only detectable at 16 h.p.i. (38). The same study found induction of both the intrinsic and extrinsic apoptotic pathways was evident only by 24 h.p.i. (42). It is clear that the transcriptional and translational profile of the host cell may differ significantly throughout the course of infection. Additionally, it must be noted that the horse (*Equus caballus*) genome is not highly annotated and thus many Ensembl gene identifications do not possess an annotated orthologue, a limiting factor in our analysis.

### What is the function of U1 and U2?

The current lack of a published reverse genetics system to study torovirus replication means we are unable to perform targeted mutagenesis. This would enable definitive experimental confirmation that U1 and U2 are translated from their respective CUG codons, followed by phenotypic analysis of knock-out mutants. However the comparative genomic analysis together with the accumulation of ribosomes on both CUG codons is highly suggestive of this being the site of initiation; CUG has previously been reported as the most commonly utilised non-AUG initiation codon in mammalian systems (43). In the case of U1, the coding sequence contains no AUG codons (in any frame), a situation that would facilitate pre-initiation ribosomes to continue scanning to the U2 CUG and the ORF1a AUG initiation sites (44). It remains a possibility that U2 translation initiates at a downstream AUG, however the only in-frame AUG is located 336 nt downstream of our presumed start site and is in a poor initiation context (‘C’ at −3) and 3’ of the ORF1a AUG. We are therefore confident that the CUG codons that were identified in the ribosome profiling data represent the genuine translational start sites.

The ORFs of both U1 and U2 are intact in all torovirus genomic sequences that we have analysed to date, including bovine (45, 46), caprine and porcine isolates (47). Most of the U2 ORF is constrained by the fact that the sequence must also retain ORF1a coding capacity in another frame. U1 is not under such limitations, although it is likely that the viral genome must maintain specific 5’ UTR structures to facilitate viral replication. Previous investigations utilising defective interfering RNAs have confirmed that no more than the first 604 nt of the 5’ UTR and the entirety of the 3’ UTR are sufficient to allow both positive and minus strand RNA synthesis (34); it is notable that this region only includes one-third of the U1 ORF (which starts at nucleotide 524) and hence only this subdomain would be constrained by maintaining two distinct functional roles. We suggest that the so-called 5’ UTR is actually limited to 523 nt preceding the CUG of U1, and the remainder of U1 and U2 is not under pressure to maintain *cis-*replication elements.

Neither ORF could be identified within the white bream virus genome, a bafinivirus that constitutes another genus within the subfamily *Torovirinae* (35), although the lack of multiple bafinivirus sequences makes comparative genomic analysis impossible.

The function(s) of the proteins encoded by both U1 and U2 remain to be elucidated. Despite the relatively large size of the U2 protein (~30 kDa), after extensive database searches no structural homologs were identified. By comparison, the U1 protein is small (~10 kDa), highly basic (pI = 10.4) and possesses many of the predicted features of a double-spanning transmembrane protein, including two hydrophobic stretches separated by a ‘hinge’ and a predicted coiled-coil tertiary topology. Based on structural similarity to known proteins, one potential function might be a virally encoded ion channel (viroporin) embedded in either intracellular or plasma membranes. It is possible that U1 plays a similar role in toroviruses to that of the coronaviral and arteriviral E proteins, which have no known toroviral homologue. The coronavirus E protein is a small transmembrane protein (~10 kDa) which possesses ion channel activity and is required for virion assembly, forming a pentamer that traverses the viral envelope (48). E proteins also possess a membrane-proximal palmitoylated cysteine residue, which is a predicted (and conserved) posttranslational modification for U1 (31).

Alternatively viroporin activity may be mediated by a small, basic double-transmembrane protein, the ORF of which is embedded within the EToV N gene in the +1 frame (with respect to N). An analogous “N+1” protein has been identified in some group II coronaviruses and is postulated to play a structural role, however it is not essential for replication (49, 50). Neither our ribosome profiling nor comparative genomic analysis provides evidence that this ORF is utilised in toroviruses. We did not observe ribosomes translating in this frame in either the initial dataset or the drug-treated samples (although Ribo-seq may not always detect poorly translated overlapping genes); further, the ORF is not preserved in all torovirus genomes.

Our data has revealed that the transcriptional landscape of a prototypic torovirus is complex and driven by many factors beyond the canonical “multi-loci TRS” model of coronaviruses. The development of a torovirus reverse genetics system would allow manipulation of potential translocation-inducing sequences and allow us to elucidate which features of the toroviral TRS cause them to act as terminators of RNA synthesis, rather than consistently inducing homology-assisted recombination. Our accompanying translational analysis has revealed two conserved novel ORFs, and has shortened the EToV 5’ UTR to a mere 523 nt. Together these data provide an insight into the molecular biology of the replication cycle of this neglected pathogen and highlight the disparities between the families of the *Nidovirales*.

## Materials and Methods

### Virus isolates

A plaque-purified isolate of equine torovirus, Berne strain (isolate P138/72) (EToV) was kindly provided by Raoul de Groot (Utrecht University) and cultured in equine dermis (ED) cells. This virus was initially isolated from a symptomatic horse in 1972 (13). ED cells were maintained in Dulbecco’s modified Eagle’s medium (Invitrogen), supplemented with 10 % foetal calf serum, 100 IU/mL penicillin, 100 µg/mL streptomycin, 1 mM non-essential amino acids, 25 mM HEPES and 1 % L-glutamine in a humidified incubator at 37°C with 5% CO_2_.

### RNA sequencing and ribosome profiling

ED cells were infected with EToV for 1 hour (h) in serum-free media (MOI = 0.1) and flash-frozen in liquid nitrogen at 8 h post infection (h.p.i.) prior to either RNA isolation or ribosome purification for profiling. Cells were either not pretreated or, where stated, were treated with a final concentration of 100 μg/mL cycloheximide (CHX) for 2 minutes (Sigma-Aldrich) or 2 μg/mL of harringtonine for 3 minutes (LKT Laboratories) followed by CHX for 2 minutes, before flash-freezing. RNA and ribosomes were harvested according to previously published protocols (15, 51) with minor modifications. Following either RPF or RNA isolation, duplex-specific nuclease was not utilised but instead rRNA was depleted with the RiboZero [human/mouse/rat] kit (Illumina). Libraries were prepared and sequenced using the NextSeq500 platform (Illumina).

### Bioinformatic analysis of Ribo-seq and RNA-seq data

Both Ribo-seq and RNA-seq reads were demultiplexed and adaptor sequences trimmed using the FASTX-Toolkit (hannonlab.cshl.edu/fastx_toolkit/). Reads shorter than 25 nt after trimming were discarded. Bowtie (version 1.2.1.1) databases were generated as follows. Horse ribosomal RNA (rRNA) sequences were downloaded from the National Center for Biotechnology Information (NCBI) Entrez Nucleotide database (accessions EU081775.1, NR_046271.1, NR_046309.2, EU554425.1, XM_014728542.1 and FN402126.1) (52). As the full-length virus RNA (vRNA) reference genome was not available for EToV, a reference was constructed from the following overlapping segments available from Entrez Nucleotide: DQ310701.1 (positions 1-14531), X52374.1 (13475–21394), X52506.1 (21250–26086), X52505.1 (26054–26850), X52375.1 (26784–27316) and D00563.1 (27264–279923). Horse messenger RNA (mRNA) sequences from EquCab2.0 (GCF_000002305.2) were downloaded from NCBI RefSeq (53). Horse non-coding RNA (ncRNA) sequences were obtained from Ensembl release 89 (54) and combined with horse transfer RNA (tRNA) sequences from GtRNADB (55). Horse genomic DNA (gDNA) was obtained from Ensembl release 89. All horse sequences were from the EquCab2.0 genome build. Trimmed reads were then mapped sequentially to the rRNA, vRNA, mRNA and ncRNA databases using bowtie version 1.2.1.1 (56), with parameters -v 2 --best (i.e. maximum 2 mismatches, report best match), with only unmapped reads passed to each following stage. Reads that did not align to any of the aforementioned databases were then mapped to the host gDNA using STAR version 2.5.4a (57), again allowing a maximum of 2 mismatches per alignment. Remaining reads were classified as unmapped.

Ribo-seq density and RNA-seq density were calculated for each gene in the EToV genome (Figure 3, Figure 8). To normalise for different library sizes, reads per million mapped reads (RPM) values were calculated using the sum of positive-sense virus RNA reads and host RefSeq mRNA reads as the denominator. In order to standardise the regions used to calculate RNA-seq and Ribo-seq density, the following regions were selected: ORF1a, start codon (position 882) to 5’ end of frameshift site (position 14518); ORF1b, 3’ end of frameshift site (position 14525) to 5’ end of the S gene hairpin (position 21118); all other ORFs, initiation codon to termination codon. For U2, a region overlapping with ORF1a was used because only 46 bases are unique to U2 and, for Figure 8, the ORF1a coordinates were updated to exclude the region which overlaps with U2, giving a range from 1552 to 21394. In addition, for all ORFs, only Ribo-seq reads mapping to the predominant phase (i.e. reads mapping to the first positions of codons) were used, as this should greatly diminish misassignment of ORF1a-translating ribosomes to U2 or *vice versa*. Reads mapping to the first five codons at the 5’ end of each region or the last six codons at the 3’ end of each region were excluded. For subgenomic RNAs, RNA-seq density was calculated for the same regions as described for Ribo-seq. For the genomic RNA the regions for ORF1a and ORF1b were combined into the interval from the start codon of ORF1a (position 882) to the 5’ end of the S gene hairpin (position 21118). Ribo-seq and RNA-seq densities were calculated as the number of reads per million mapped reads for which the 5’ end maps to each region, divided by the length of the region in nt, multiplied by 1000 (i.e. RPKM). For RNA-seq, a decumulation strategy was used to subtract the estimated RNA-seq density for longer overlapping genomic and subgenomic transcripts that would contribute to the RNA-seq density measured for each of the 3’ ORFs: the genomic RNA-seq density was subtracted from all subgenomic densities, and then the RNA-seq densities of overlapping “upstream” subgenomic transcripts were iteratively subtracted from “downstream” regions (e.g. RNA-seq density in the unique region of M was subtracted from HE, and this was subtracted from N). Translation efficiency for each gene was calculated as Ribo-seq density / decumulated RNA-seq density. Translational efficiencies for HE could not be accurately estimated as the low expression of the HE transcript made the decumulation procedure for HE susceptible to noise.

Read length distributions were calculated for Ribo-seq and RNA-seq reads mapping to positive-sense host mRNA annotated CDSs or to the positive- or negative-sense EToV genome (Supplementary Figure 1). Histograms of host mRNA Ribo-seq and RNA-seq 5’ end positions relative to initiation and termination codons (Supplementary Figure 2, Supplementary Figure 3) were derived from reads mapping to mRNAs with annotated CDSs ≥ 450 nt in length and annotated 5’ and 3’ UTRs ≥ 60 nt in length. Host mRNA Ribo-seq and RNA-seq phasing distributions (Supplementary Figure 4) were calculated taking into account interior regions of annotated coding ORFs only (specifically, reads for which the 5’ end mapped between the first nucleotide of the initiation codon and 30 nt 5’ of the termination codon) in order to exclude reads on or near initiation or termination codons. For viral genome coverage plots, but not for meta-analyses of host RefSeq mRNA coverage, mapping positions of RPF 5’ ends were offset + 12 nt to approximate the location of the ribosomal P-site (15).

### Analysis of viral transcripts

The EToV (Berne isolate) genome sequence was confirmed by *de novo* assembly of unmapped and vRNA reads from the infected RNA-seq samples. Assembly was performed using Trinity (58) with the default settings for stranded single ended (--SS_lib type “F”) data. Viral contigs were identified using BLASTN (27) against a database of EToV reference sequences based on the NCBI records listed above. The viral contig was aligned to the reference using the MAFFT L-INS-i method (59).

Chimaeric reads were classified as reads for which the entire read mapped uniquely to the viral genome, with no mismatches, after adding a single breakpoint, with a minimum of 12 nt mapping on either side of the breakpoint, at least 5 nt apart. To identify such reads, all unmapped reads were split into two sub-reads at every possible position ≥12 nt from either end and these sub-reads were mapped to the viral genome using bowtie with no mismatches and no multimapping permitted. Transcription junctions were defined as “donor/acceptor” pairs that were either supported by at least 10 chimaeric reads or contained the entire 5’ leader and TRS sequence in the 5’ segment of the read. At some positions single nucleotide resolution for the chimaeric break-point could not be established; where reads were found to break at adjacent possible positions these positions were merged to give a short region containing the breakpoint. The number of non-chimaeric reads spanning each donor and acceptor site was calculated as the number of reads which overlapped the site by at least 12 nt in either direction (as chimaeric reads overlapping the site by < 12 nt are not detectable). The proportion of chimaeric reads at each “donor” or “acceptor” site is therefore the number of chimaeric reads with a breakpoint at the site divided by this number plus the number of non-chimaeric reads spanning the site (Figure 4B).

To visualise TRS conservation, multiple sequence alignments were generated using Clustal Omega with default parameters (60). RNA structure was predicted using RNA-Alifold (61) and visualised using VARNA (62).

### Differential gene expression analysis

For analysis of host differential expression between non-drug treated infected and mock-infected cells, all reads which did not map to rRNA or vRNA were mapped to the EquCab2.0 reference genome and annotations (Ensembl release 89) using STAR (57) with a maximum of two mismatches and removal of non-canonical, non-annotated splice junctions. Read counts were generated using HTSeq 0.8.0 (63). For differential transcription analysis, gene level counts were generated across the Ensembl release 89 EquCab2.0 gtf file, filtered to include only protein-coding genes. For differential translation efficiency analysis only coding regions (CDS) were considered: both RNA-seq and Ribo-seq counts were generated at CDS level using intersection-strict mode, based on the same annotation set. Multimapping reads were excluded from both analyses. Differential transcript abundance analysis was performed using the standard DESeq2 (64) pipeline described in the vignette. Genes to which <10 reads mapped were discarded and shrinkage of log_2_ fold changes for lowly expressed genes was performed using the lfcshrink method of DESeq2. All recommended quality control plots were inspected, and no major biases were identified in the data. False discovery rate (FDR) values were calculated using the R fdrtool package (65). Genes with a log_2_ fold change >1 and an FDR less than 0.1 were considered to be differentially expressed. Gene ontology (GO) term enrichment analysis (66) was performed against a background of all horse protein-coding genes in the Ensembl gtf using a Fisher Exact Test and corrected for multiple testing with a Bonferroni correction. GO annotations for horse genes were downloaded from BiomaRt (Ensembl release 90) (67). Differential translational efficiency analysis was carried out using the CDS counts table, normalised using the DESeq2 “sizeFactors” technique. Similar to the differential transcription analysis, genes to which <10 reads mapped were discarded. Again all recommended quality control plots for DESeq2 were inspected and no major biases were identified in the data. Differential translation efficiency analysis was performed using Xtail (68), following the standard pipeline described in the vignette. *P*-values were adjusted automatically within Xtail using the Benjamini–Hochberg method. Genes with a log_2_ fold change >1 and an adjusted *p*-value less than 0.1 were considered to be differentially translated. GO enrichment analysis was performed as described for the differential transcript abundance analysis.

### Comparative genomics

The Genbank accession numbers utilised for comparative genomic analysis were as follows: DQ310701.1 (Berne virus), AY427798.1 (Breda virus) (45), KR527150.1 (goat torovirus), JQ860350.1 (porcine torovirus) (47), KM403390.1 (porcine torovirus) (69), LT900503.1 (porcine torovirus), LC088094.1 (bovine torovirus) and LC088095.1 (bovine torovirus) (46). The ratio of nonsynonymous to synonymous substitution rates (dN/dS) was estimated using the codeml program in the PAML package (22). The eight torovirus U1 nucleotide sequences were translated, aligned as amino acids with MUSCLE (70), and the amino acid alignment used to guide a codon-based nucleotide alignment (EMBOSS tranalign) (71). Alignment columns with gap characters in any sequence were removed, resulting in a reduction from 81 to 79 codon positions. PhyML (72) was used to produce a nucleotide phylogenetic tree for the U1 alignment and, using this tree topology, dN/dS was calculated with codeml. The standard deviation for the codeml dN/dS value was estimated via a bootstrapping procedure, in which codon columns of the alignment were randomly resampled (with replacement); 100 randomized alignments were generated, and their dN/dS values calculated with codeml.

Coding potential within each reading frame was analysed using MLOGD (23) and synonymous site conservation was analysed with synplot2 (29). For these analyses we generated a codon-respecting alignment of the eight torovirus full-genome sequences using a procedure described previously (29). In brief, each individual genome sequence was aligned to a reference sequence using code2aln version 1.2 (73). Breda virus (GenBank accession AY427798) was used as reference, since unlike Berne virus it contains an intact HE gene. Genomes were then mapped to reference sequence coordinates by removing alignment positions that contained a gap character in the reference sequence, and these pairwise alignments were combined to give the multiple sequence alignment. This was analysed with MLOGD using a 40-codon sliding window and a 5-codon step size. For each of the three reading frames, within each window the null model is that the sequence is non-coding whereas the alternative model is that the sequence is coding in the given reading frame. Positive/negative values indicate that the sequences in the alignment are likely/unlikely to be coding in the given reading frame. To assess conservation at synonymous sites, the concatenated coding regions were extracted from the alignment and analysed with synplot2.

## Data availability

The sequencing data reported in this paper have been deposited in ArrayExpress (http://www.ebi.ac.uk/arrayexpress) under the accession number E-MTAB-6656.

## Acknowledgements

We thank Raoul de Groot and Arno van Vliet (Utrecht University) for providing the virus isolates and helpful advice, and Polly Roy (London School of Hygiene and Tropical Medicine) for ED cells.

---

**Supplementary Table 1.** Read counts for each sample. Too short, adaptor only, rRNA forward/reverse, mRNA forward/reverse, ncRNA forward/reverse, gDNA forward/reverse, and vRNA forward/reverse reads were summed to give the total mapped read count. Remaining reads were classified as unmapped.

**Supplementary Table 2.** Gene descriptions (Ensembl gene identifiers and gene symbols) for transcripts which were differentially transcribed or translated, in EToV compared to mock infected cells.

**Supplementary Figure 1.**
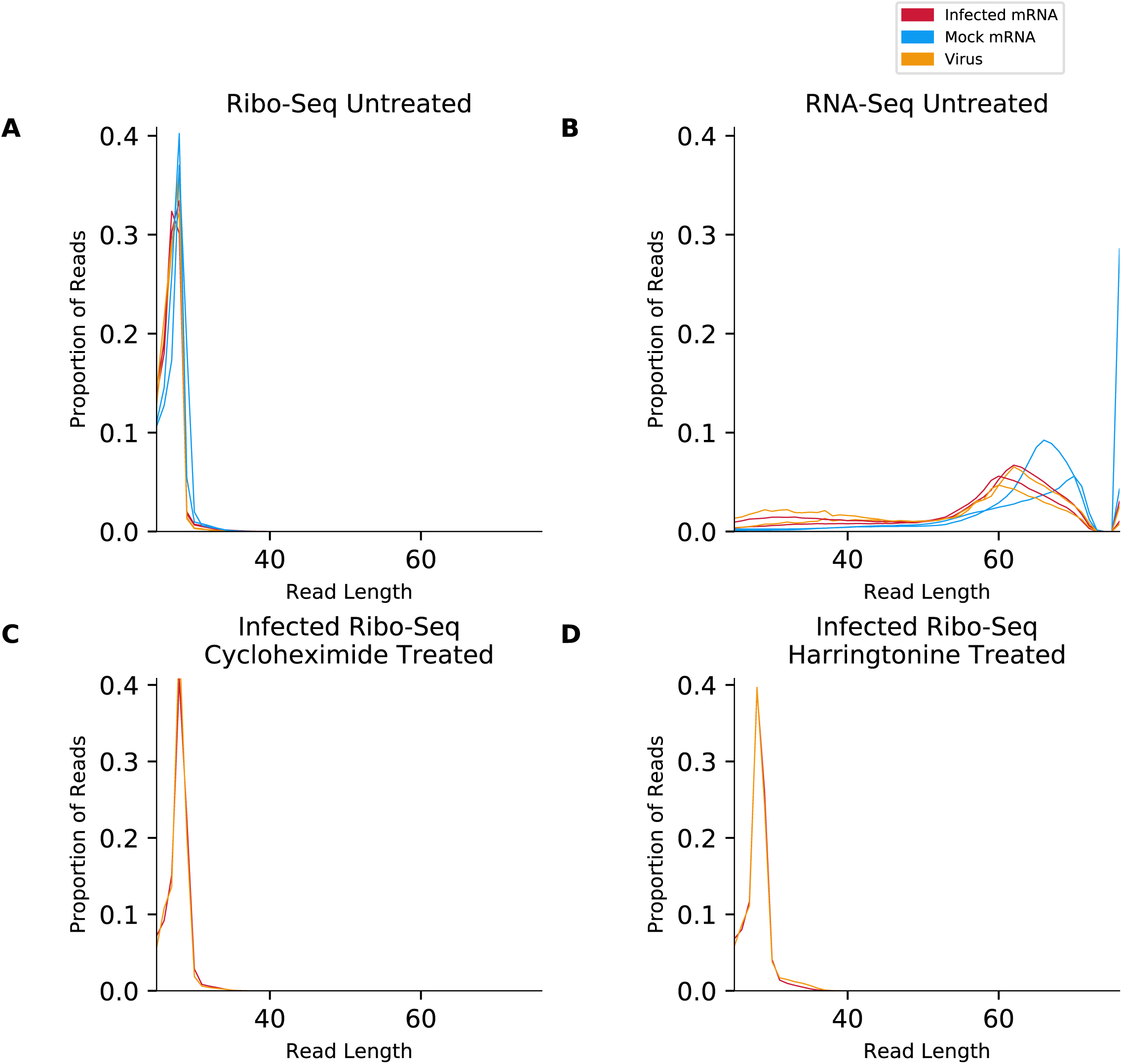
Comparison of read length distribution for reads mapping to EToV in infected cells (orange), host mRNAs in non-infected cells (blue) and host mRNAs in infected cells (red) for (A) Ribo-seq data in non-drug treated cells; (B) RNA-seq data in non-drug treated cells; (C) Ribo-seq data in cycloheximide-treated cells; and (D) Ribo-seq data in harringtonine-treated cells.

**Supplementary Figure 2.**
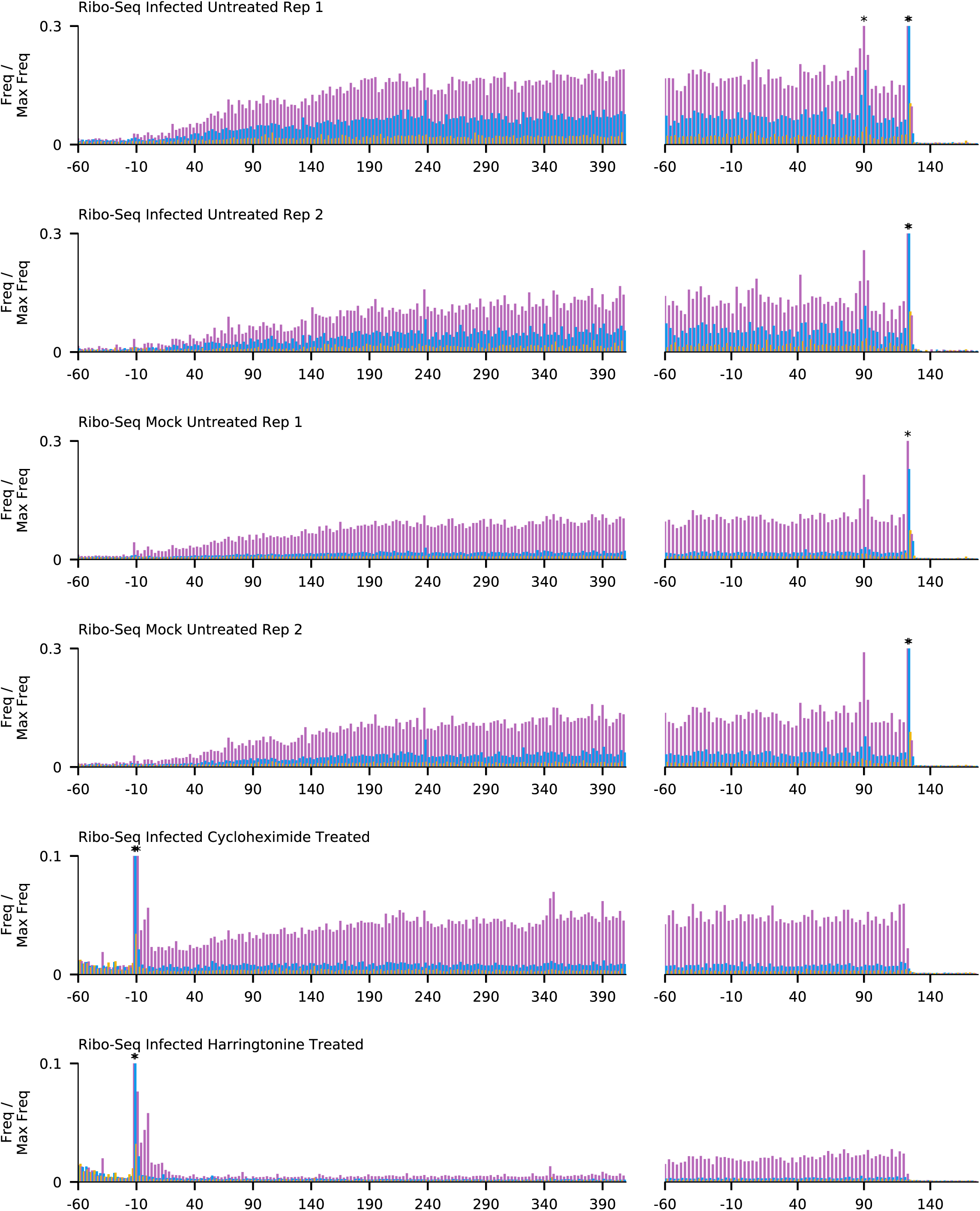
Histograms of Ribo-seq read 5’ end positions (nt) relative to annotated initiation (left) and termination (right) sites, summed across all host mRNAs. Bars are coloured by phase relative to the first base of the start codon (pink: phase 0; blue: phase +1; yellow: phase −1). Histograms are scaled so that the maximum value is 1. For clarity, the y-axis is cropped at 0.3 for non-drug treated and 0.1 for drug treated cells; bars which extended beyond this point are marked with an asterisk (*).

**Supplementary Figure 3.**
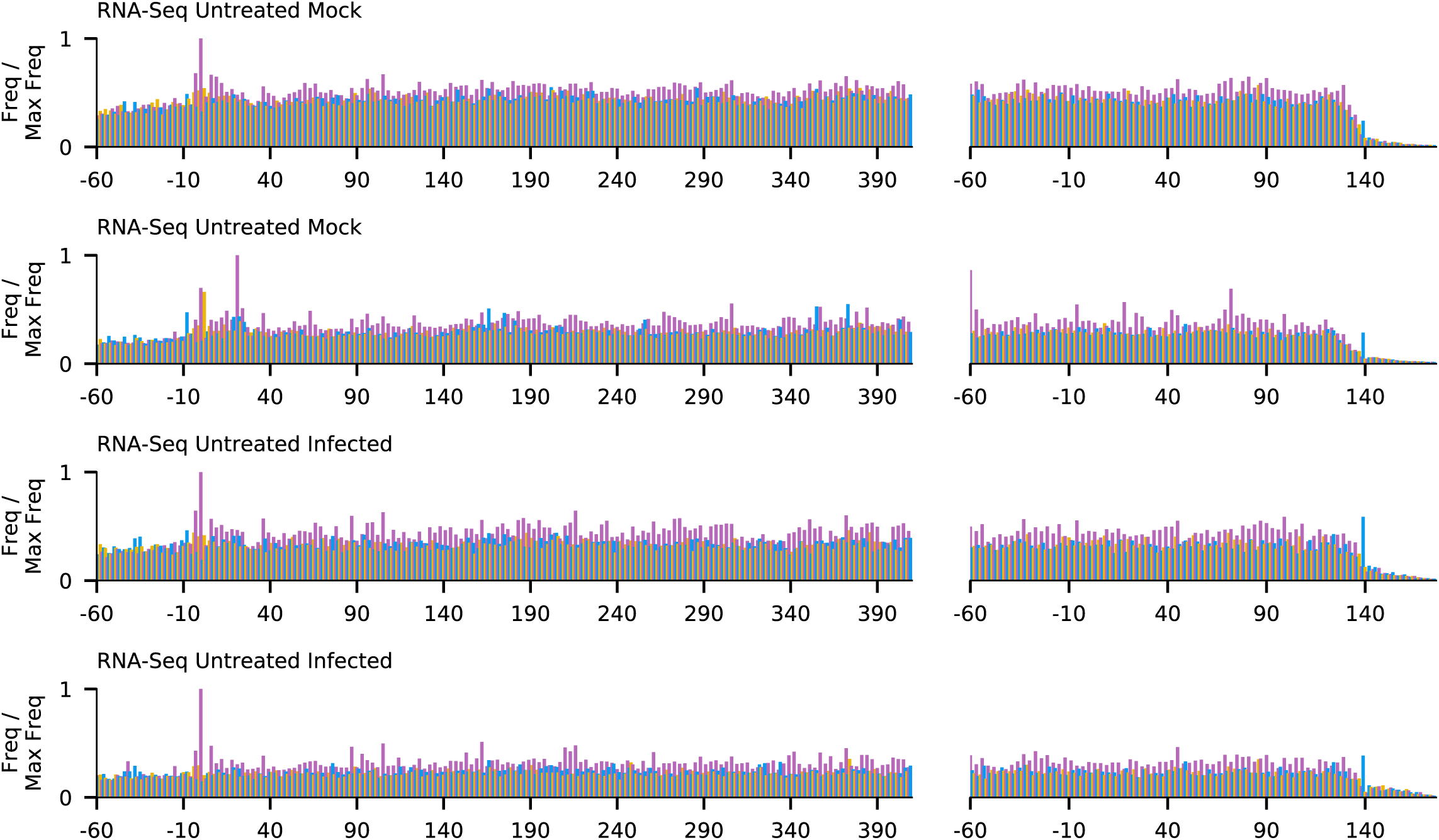
Histograms of RNA-seq read 5’ end positions (nt) relative to annotated initiation (left) and termination (right) sites, summed across all host mRNAs. Bars are coloured by phase relative to the first base of the start codon (pink: phase 0; blue: phase +1; yellow: phase −1). Histograms are scaled so that the maximum value is 1.

**Supplementary Figure 4.**
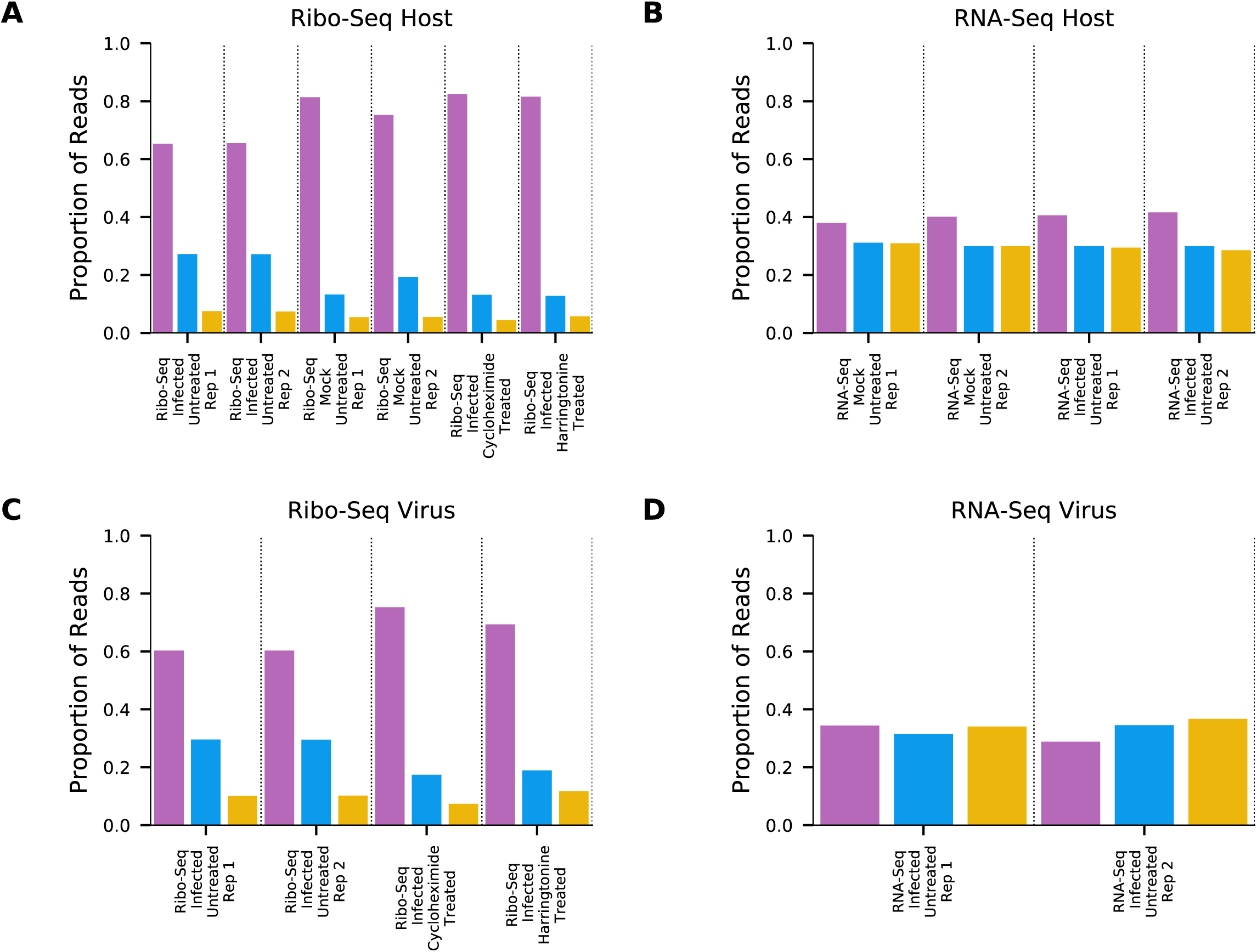
Phasing of the 5’ ends of reads (pink: phase 0; blue: phase +1; yellow: phase −1) for (A) Ribo-seq reads mapping to host mRNA coding regions, (B) RNA-seq reads mapping to host mRNA coding regions, (C) Ribo-seq reads mapping to virus mRNA coding regions and (D) RNA-seq reads mapping to virus mRNA coding regions.

**Supplementary Figure 5.**
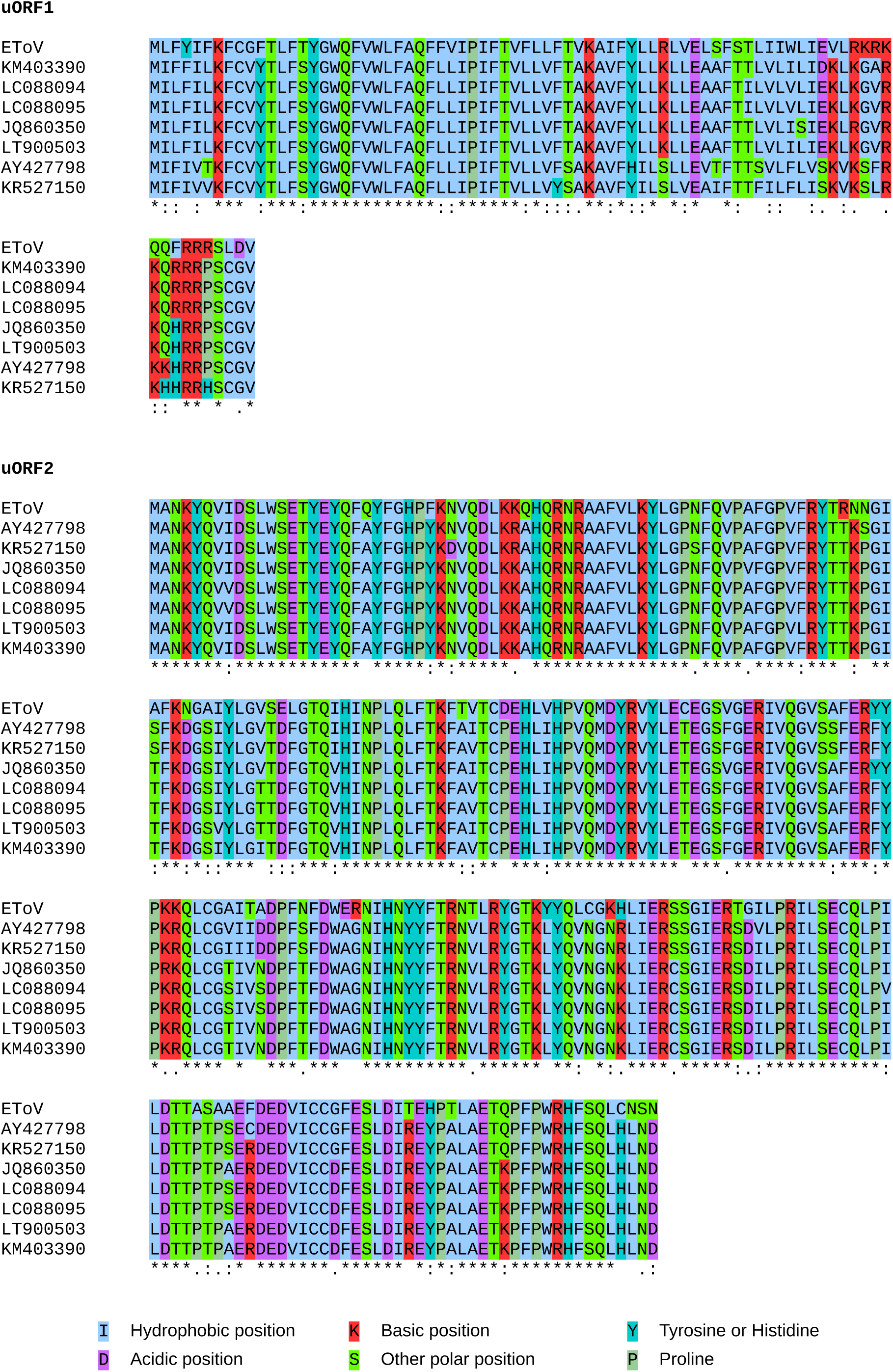
Conservation of uORF1 and uORF2 in the eight publicly available torovirus genomes. Individual amino acid residues are coloured according to their biochemical properties.

**Supplementary Figure 6.**
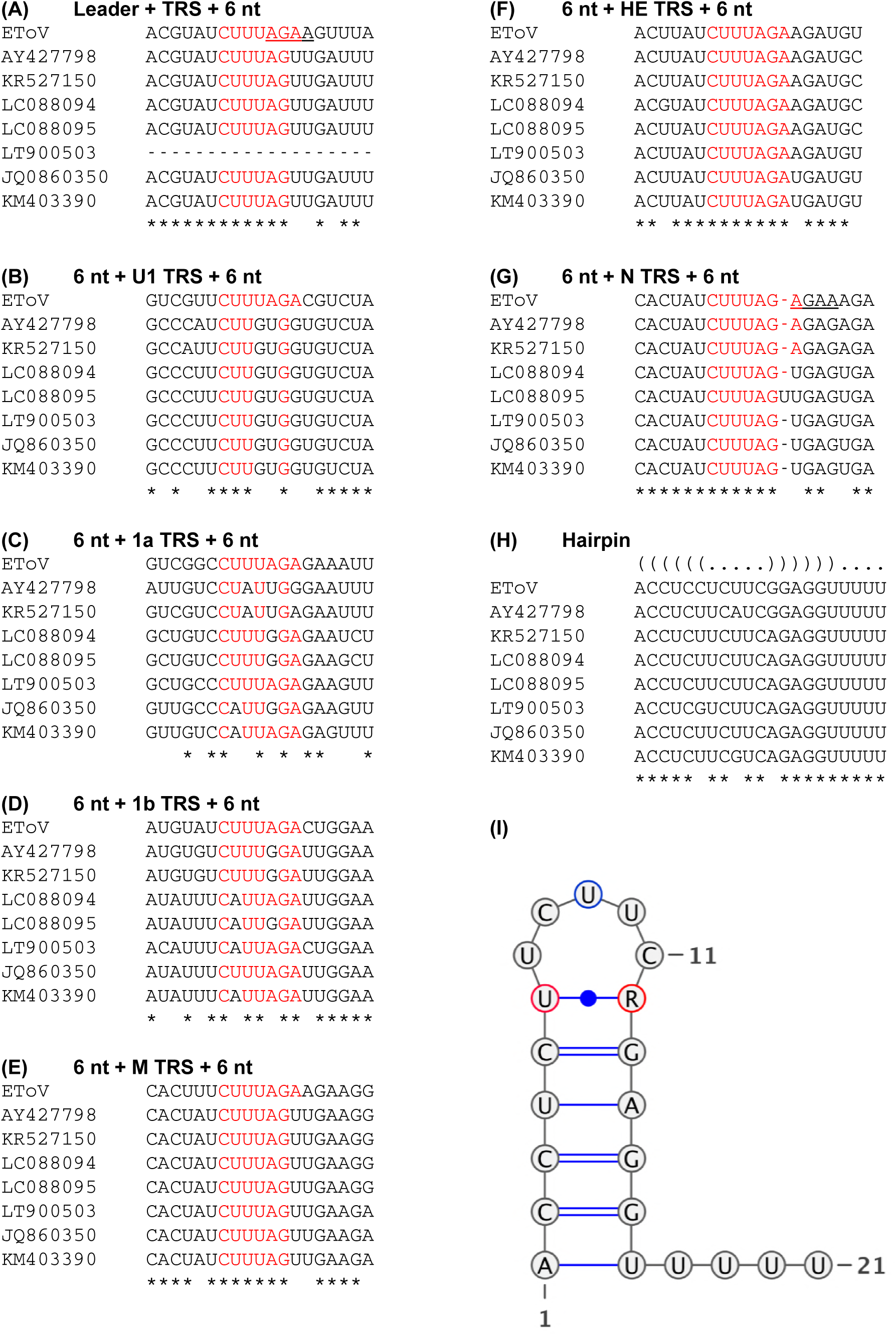
Conservation of TRSs and regulatory structures in the eight publicly available torovirus genomes. Regions were selected based on the presence of a putative TRS in the EToV genome. The TRS and six flanking nucleotides are displayed; putative TRS nucleotides are highlighted in red. Nucleotide conservation between all eight sequences is indicated by an asterisk (*). The predicted hairpin structure (I) is based upon nucleotide conservation across all eight genomes. Variant nucleotides are circled in either red (covariance indicates the predicted pairing may occur in all but one genome) or blue (variable). R indicates a purine exists in all genomes.

